# Kidney cystogenesis in embryonic- and adult-onset ADPKD is suppressed from lack of adenylyl cyclase targeting to cilia

**DOI:** 10.1101/2025.05.27.656198

**Authors:** Sun-hee Hwang, Kyungsuk Choi, Hemant Badgandi, Kevin White, Yu Xun, Feng Qian, Saikat Mukhopadhyay

## Abstract

Multiple cellular pathways are dysregulated in autosomal dominant polycystic kidney disease (ADPKD), but mechanisms initiating cyst formation are unknown. ADPKD is caused by mutations in *Pkd1/Pkd2* genes that encode for polycystins that localize to primary cilia. The primary cilium is a miniscule subcellular compartment for generating signaling outputs that profoundly affect cellular function. Severe cystogenesis from polycystin loss is mostly cilia dependent. However, the polycystin-repressed ciliary signals that promote cyst growth are unknown and have been challenging to uncouple from downstream cystogenic pathways. Here we aimed at differentiating ciliary adenylyl cyclase signaling from total cellular changes in second messenger cAMP implicated in cystogenesis. We studied an Ankyrin repeat and MYND domain protein, ANKMY2 that we previously implicated in maturation and ciliary localization of adenylyl cyclases in fibroblasts. We studied kidney-specific conditional knockout mouse models of *Ankmy2/Pkd1* and ciliary localization of adenylyl cyclases in kidney epithelial cells. We found suppression of early postnatal renal cystogenesis and prolonged survival in an embryonic onset *Pkd1* deletion model from ANKMY2 loss. Phosphorylated CREB formation, from elevated cellular cAMP levels, remained unaffected. Cyst load in male mice in an adult inducible conditional *Pkd1* deletion model was suppressed from ANKMY2 loss. Mechanistically, ANKMY2 determined ciliary trafficking of adenylyl cyclases in kidney epithelial cells without disrupting cilia. Further, ANKMY2 loss prevented ciliary length increase in ADPKD mouse models irrespective of cyst load or sex. Cilia length increase was seen preceding cystogenesis. Our results suggest that targeting of adenylyl cyclases to renal epithelial cilia promotes PC1/2-inhibited cilia-dependent cyst activation distinct from cyst progression involving cellular cAMP.

## Introduction

The kidney nephron tubular epithelial cells have primary cilia starting from development and continuing into adult life.^1^ The primary cilium is a paradigmatic subcellular compartment for generating signaling outputs that can have profound effects on cellular function.^2^ The role of primary cilia in kidney tubule morphogenesis and in the related disease such as polycystic kidney disease is complex^3, 4^ and is also temporally regulated.^5^ Based on genetic epistasis performed using cilia loss in the background of polycystic kidney disease, both positive and negative counterregulatory signals generated by the cilia has been proposed to regulate severity of cystogenesis.^3, 4^ Furthermore, temporal postnatal developmental windows of ciliary function have been shown to determine the timing of cyst initiation, with cilia loss before P14 driving early cystogenesis.^5^ Cyst progression is regulated by a multifactorial and multi-organellar process,^6^ thus studying the precise contribution of cilia in cystogenesis has been complicated. Furthermore, cilia localized proteins have extraciliary components and teasing apart ciliary from extraciliary function from knockout studies is not feasible.^7^ Lack of cilia prevents studying signals that are generated by cilia, therefore studying ciliary function by preventing ciliary compartmentalization without disrupting cilia is needed to study ciliary contributions in downstream pathways.

ADPKD is caused by mutations in genes (*Pkd1* and *Pkd2*) encoding for polycystin-1 (PC1) and polycystin-2 (PC2)—both of which localize to primary cilia.^8, 9^ Loss of polycystins causes severe cystogenesis, which is mostly cilia dependent, suggesting that such loss derepresses cyst activators.^10^ However, the polycystin-inhibited cilia-dependent/localized cyst activator(s) (CDCA) that promote cyst growth in ADPKD are unknown. The CDCA should in principle localize to cilia, should not affect cystogenesis on being depleted but should prevent cystogenesis in embryonic- or adult-onset cystogenesis upon concomitant loss in the background of PC1/2 depletion.^4^ A clue to the positive signal(s) came from studying TULP3, a ciliary trafficking adapter pivotal in ciliary membrane composition that functions in coordination with the IFT-A complex.^11^ Whereas deletion of *Tulp3* and IFT-A subunit *Ift139* exacerbates early postnatal cystogenesis from *Pkd1/2* loss,^12, 13^ similar deletion later in adult-onset disease causes inhibition of cystogenesis.^12, 14^ The differences in genetic epistasis between *Tulp3* and *Pkd1* between early- and adult-onset disease^4^ is explained by TULP3 and IFT-A determining trafficking of multiple cargoes to the ciliary membrane.^11^ For example, the exacerbation of early-onset disease from TULP3 loss is at least partly to be likely from lack of trafficking of ARL13B, a bonafide TULP3 cargo, to cilia.^15–17^ The ciliary components of CDCA signal(s) in ADPKD cystogenesis remain unknown, although cytoplasmic components downstream of cilia, such as cell cycle kinases^18^, the transcription factor GLIS2,^19^ and the ankyrin-repeat protein ANKS3,^20^ have recently been described.

Previous studies have demonstrated that cellular cAMP is a major driver of cystogenesis in PKD^21^ and in other cystic kidney disease models including ARPKD^22^, NPHP^23^ and juvenile cystic kidney disease such as from loss of NEK8.^24^ Stimulation by cAMP analogues such as 8-Br-cAMP has also been shown to contribute to cyst growth by stimulating fluid secretion^25^ through activation of the CFTR chloride channel^26, 27^ and by increasing cell proliferation through activation of the B-Raf/MEK/ERK pathway.^28^ Conversely, drugs that reduce intracellular cAMP levels, such as vasopressin receptor 2 (V2R) antagonists and somatostatin receptor agonists, inhibit cyst growth in ADPKD^29^ and other cystic kidney disease mice models.^23^ Together, these drugs reduce cellular cAMP levels, reduce cystic burden in animal models^30^ and have renoprotective effects in ADPKD patients.^31^ Ciliary volume is less than 1/30000 of the total cellular volume.^32^ Because of miniscule volume of cilia compared to the cytoplasm, any changes in ciliary cAMP are unlikely to cause global changes in cellular cAMP.^33, 34^ Levels of cAMP that are elevated in cyst epithelial cells from humans with ADPKD and in cystic kidneys from *Pkd1* and *Pkd2* mutant mice are therefore not likely directly from diffusion of ciliary cAMP into the cytoplasm.

Membrane adenylyl cyclases are topologically complex enzymes having two transmembrane bundles of 6 transmembrane domains each that bring together two pseudosymmetric cyclase domains catalyzing production of cAMP from ATP.^35^ At least three of the nine membrane adenylyl cyclases traffic to cilia, including ADCY3, ADCY5, and ADCY6.^36–38^ The role of membrane adenylyl cyclases ADCY5 and ADCY6 were previously tested separately in PKD models using straight knockout^39^ or conditional knockout strategies^40^, respectively, which were shown to have partial amelioration of cystic burden. Knockouts or conditional knockouts of *Adcy5* and *Adcy6* also affect total cellular cAMP levels.^39, 40^ ADCY3 has also been shown to be expressed in ADPKD kidney cysts.^41^ Downregulation of the cAMP effector protein kinase A (PKA) using kidney-specific expression of a dominant negative PKA regulatory subunit *RIαB* allele delays development and progression of PKD in *Pkd1*^RC/RC^ models.^42^ The promotion of cystogenesis by PKA in ADPKD mice models is also through modulation of total cellular cAMP levels.^42^ Correspondingly, kidney-specific knockout of PKA regulatory subunit *RIα* upregulates PKA activity, induces cystic disease in wild-type mice and aggravates it in *Pkd1*^RC/RC^ mice.^43^ Whereas, cAMP levels in these models increase with the severity of cystic disease,^44^ these studies do not account for the ciliary component of cAMP signaling in pathogenesis of ADPKD.

Some membrane adenylyl cyclases have been shown to be bona fide IFT-A cargoes to cilia.^45^ However, dissecting specific role of ciliary compartmentalization of multiple adenylyl cyclases while still retaining cellular pools and without disrupting cilia has been challenging. Here we take an approach to selectively study adenylyl cyclase signaling from cilia in both embryonic-onset and adult-onset mouse ADPKD models. We use nephron-specific conditional knockouts of a recently described Ankyrin repeat and MYND domain containing protein ANKMY2 that regulates trafficking of multiple adenylyl cyclases to cilia without totally depleting cellular levels,^46^ unlike adenylyl cyclase knockouts.^39, 40^ We find suppression of early cystogenesis in both embryonic-onset and adult-onset disease from loss of targeting of adenylyl cyclases to cilia by ANKMY2. We also find suppression of ciliary length increase in *Pkd1* deleted kidney epithelial cells from ANKMY2 loss. Furthermore, we find that cilia length increase precedes cystogenesis. These results suggest that ANKMY2-mediated ciliary cAMP signaling promotes cilia-dependent cyst activation from loss of polycystins. Our studies on ciliary drivers of cystogenesis and temporal windows of ciliary trafficking regulation will be fundamentally important in devising ADPKD treatment strategies.

## Results

### Early cystogenesis in embryonic-onset PKD is suppressed from lack of ANKMY2

We previously demonstrated high hedgehog signaling in the mouse embryonic neural tube in the *Ankmy2* knockout that was fully cilia-dependent from loss of targeting of multiple adenylyl cyclases to cilia.^46^ Previous studies have tested the role of ADCY5 and ADCY6 separately in PKD models using straight knockout^39^ or conditional knockout strategies,^40^ respectively. Lack of ANKMY2 allowed us to test the specific role of multiple adenylyl cyclases without generating knockouts of multiple adenylyl cyclase while still retaining cellular pools. However, as *Ankmy2* knockouts are E8.5 lethal, we generated a conditional knockout allele of *Ankmy2.*^46^ Using the collecting duct-specific *HoxB7-Cre* that starts expressing in the embryonic kidney, we generated conditional knockouts of *Ankmy2* along with *Pkd1.* By P3, the *HoxB7-Cre*; *Pkd1^f/f^* animals had significantly increased 2-kidney/body weight and kidney cystic index. Notably, *HoxB7-Cre*; *Pkd1^f/f^*; *Ankmy2^f/f^* animals showed a significant reduction in kidney to body weight and cystic index compared to *HoxB7-Cre*; *Pkd1^f/f^*animals (**Figure 1A-C**). Hematoxylin and Eosin (H&E) staining for all kidneys assessed are shown in **Supplemental Figure 1A**. Immunostaining of kidney sections confirmed cysts in AQP2+ collecting ducts in *HoxB7-Cre*; *Pkd1^f/f^*that was reversed in *HoxB7-Cre*; *Pkd1^f/f;^ Ankmy2^f/f^* mice (**Figure 1D**). The *HoxB7-Cre*; *Ankmy2^f/f^*animals were not affected (**Figure 1A-E** and **Supplemental Figure 1A**). Transcript levels of *Pkd1* and *Ankmy2* collected from total kidneys trended towards being reduced in corresponding conditional knockouts, respectively, compared to controls (**Supplemental Figure 1B**). As epithelial cells in nephrons at this early postnatal stage are still proliferating, we tested transcript levels of *Lipocalin-2* (*LCN2*) as an orthogonal measure of cystogeneisis. LCN2, also known as neutrophil gelatinase-associated lipocalin (NGAL), is associated with cyst expansion in PKD in rodent models and humans.^47^ Interestingly, levels of *Lcn2* transcripts significantly increased in *Hoxb7-Cre*; *Pkd1^f/f^* cystic kidneys compared to *Hoxb7-Cre*; *Pkd1^f/f^*; *Ankmy2^f/f^*P3 kidneys (**Figure 1F**). Thus, early cystogenesis in embryonic-onset PKD is suppressed from lack of ANKMY2.

**Figure 1.**
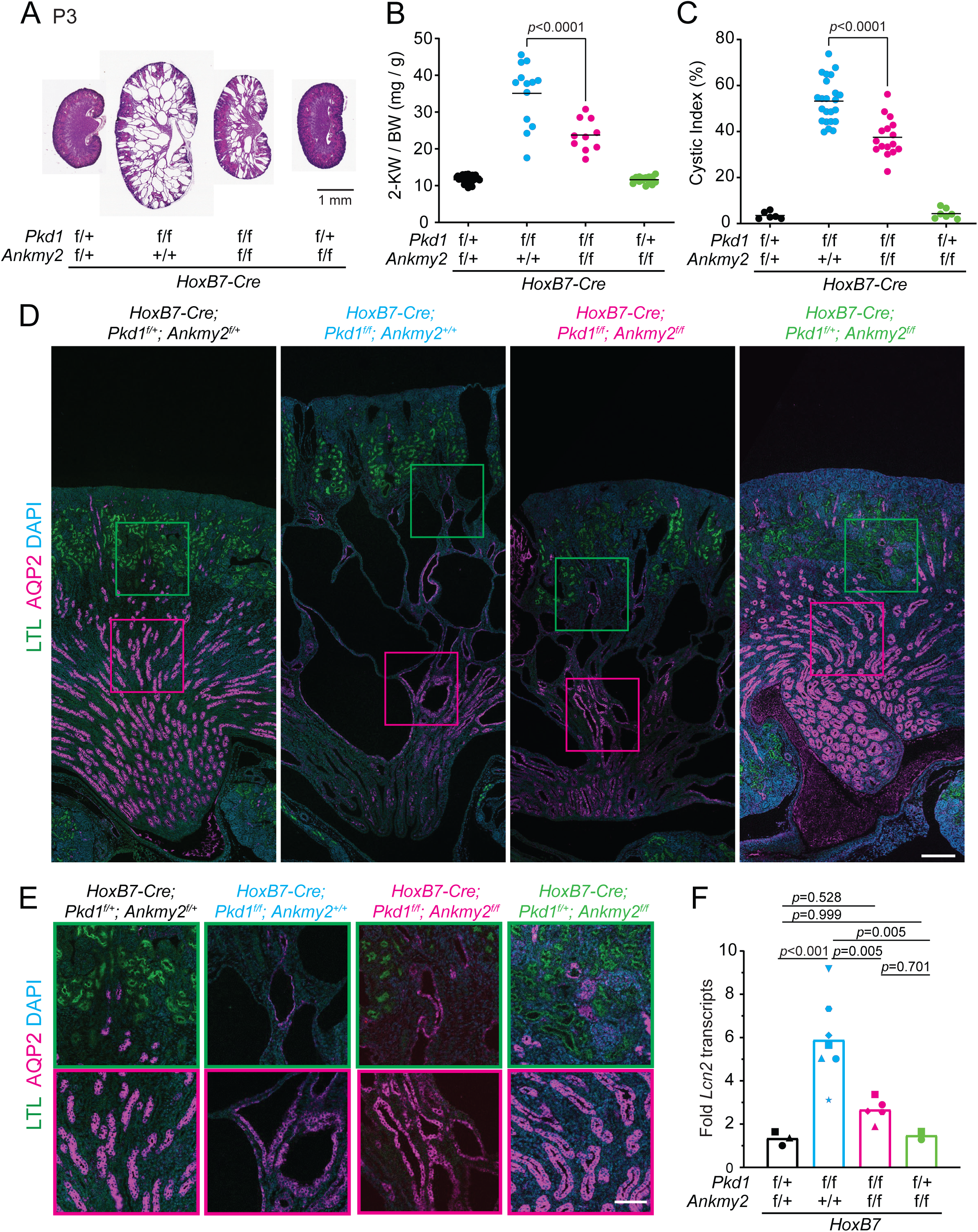
Early cystogenesis in embryonic-onset PKD is suppressed from lack of ANKMY2. **(A)** H&E images of kidneys in control, *HoxB7-Cre; Pkd1^f/f^*, *HoxB7-Cre; Pkd1^f/f^; Ankmy2^f/f^*, *HoxB7-Cre; Ankmy2^f/f^*mice at P3 in C57BL/6J background. Images of multiple kidneys shown in Figure S2B. **(B)** 2-Kidney to body weight ratios at P3 in *HoxB7-Cre; Pkd1^f/f^* mice were significantly higher than *HoxB7-Cre; Pkd1^f/f^; Ankmy2^f/^*. P3 animals had body weight between 2.5-3 g. **(C)** Significantly increased cystic index in *HoxB7-Cre; Pkd1^f/f^* mice compared to *HoxB7-Cre; Pkd1^f/f^; Ankmy2^f/f^*. **(D-E)** AQP2/LTL co-staining in kidney sections at P3 shows multiple Aqp2 positive cysts in medullar and corticomedullar junctions in *HoxB7-Cre; Pkd1^f/f^* animals compared to *HoxB7-Cre; Pkd1^f/f^; Ankmy2^f/f^*. Scale: 200 µm. Magnified images are shown in E. LTL positive, green outlines; Aqp2 positive, magenta outlines. Scale: 200 µm (D), 100 µm (E). **(F)** Transcript levels of *Lnc2* in whole kidneys from *HoxB7-Cre; Pkd1^f/f^* mice are significantly higher than *HoxB7-Cre; Pkd1^f/f^; Ankmy2^f/f^*. See also Supplemental Figure 1.

### Lack of ANKMY2 improves life expectancy in embryonic-onset PKD despite having no effect on later cystic burden or cellular cAMP response

As early postnatal cystogenesis in *HoxB7-Cre*; *Pkd1^f/f^*was ANKMY2-dependent, we next checked the life expectancy and progressive cystic burden of these mice. Interestingly, the life expectancy of *HoxB7-Cre*; *Pkd1^f/f^* animals was significantly less compared to *HoxB7-Cre*; *Pkd1^f/f^*; *Ankmy2^f/f^*animals, whereas *HoxB7-Cre*; *Ankmy2^f/f^* animals remained unaffected similar to controls (**Figure 2A**). However, at P14-P15, the 2-kidney to body weight and cystic index of the kidneys were not significantly different between *HoxB7-Cre*; *Pkd1^f/f^* and *HoxB7-Cre*; *Pkd1^f/f;^ Ankmy2^f/f^* animals irrespective of sex (**Figure 2B-D**). We next tested if cAMP-dependent phosphorylation of CREB is affected from loss of ANKMY2. We noted robust nuclear localization of pCREB by immunofluorescence in *HoxB7-Cre*; *Pkd1^f/f^* mice in later postnatal kidneys at P15. The nuclear localization of pCREB at P15 was also similar between *Hoxb7-Cre*; *Pkd1^f/f^*; *Ankmy2^f/f^* and *Hoxb7-Cr*e; *Pkd1^f/f^* animals (**Figure 2E**). H&E staining for all kidneys assessed are shown in **Supplemental Figure 2.** The levels of pCREB in nucleus are reflective of cellular cAMP levels that activates protein kinase A.^48, 49^ These results suggested that total cellular cAMP level responses remained unaffected from *Ankmy2* deletion at this later stage with similar cystic burden. Therefore, lack of ANKMY2 suppressed early cystogenesis in embryonic-onset PKD and improved life expectancy with no effect on later cystic burden.

**Figure 2.**
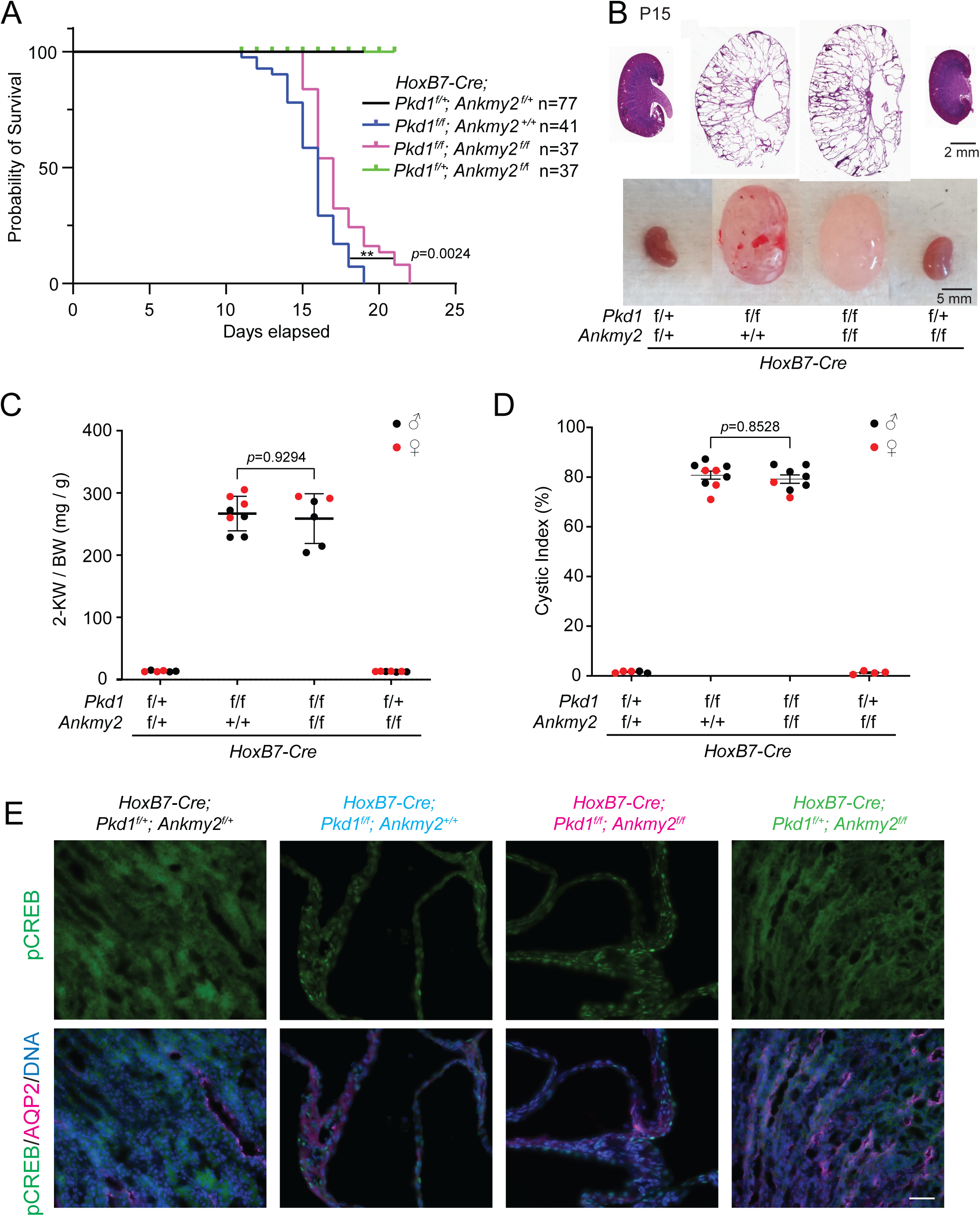
Lack of ANKMY2 improves life expectancy in embryonic-onset PKD despite having no effect on later cystic burden. (**A**) Kaplan-Meier survival curves of control (N=77), *HoxB7-Cre; Pkd1^f/f^* (N=41), *HoxB7-Cre; Pkd1^f/f^; Ankmy2^f/f^* (N=37) and *HoxB7-Cre; Ankmy2^f/f^*(N=37) mice. **, p=0024 by log-rank (Mantel-Cox) test. **(B)** H&E and whole mount images of P15 kidneys in control, *HoxB7-Cre; Pkd1^f/f^*, *HoxB7-Cre; Pkd1^f/f^; Ankmy2^f/f^*, *HoxB7-Cre; Ankmy2^f/f^*mice at P15 in C57BL/6J background. Images of multiple kidneys shown in Figure S3. **(C)** 2-Kidney to body weight ratios at P15 in *HoxB7-Cre; Pkd1^f/f^* mice were not significantly different from *HoxB7-Cre; Pkd1^f/f^; Ankmy2^f/f^*. Both males (black) and females (red) are shown. **(D)** At P15 cystic index in *HoxB7-Cre; Pkd1^f/f^* kidneys compared to *HoxB7-Cre; Pkd1^f/f^; Ankmy2^f/f^* were not significantly different. **(E)** AQP2/pCREB co-staining in kidney sections at P15 shows comparable nuclear pCREB levels in *HoxB7-Cre; Pkd1^f/f^* animals and *HoxB7-Cre; Pkd1^f/f^; Ankmy2^f/f^*cysts. Scale: 50 µm. See also Supplemental Figure 2.

### Cystogenesis in adult-onset PKD is suppressed from lack of ANKMY2

We next tested the role of ANKMY2 in adult-onset PKD models. We utilized the pan nephron-specific deletion using the doxycycline inducible *Pax8^rtTA^*; *TetO-Cre* system to delete *Pkd1* and/or *Ankmy2* using conditional alleles. We injected Doxycycline at P28-30 and performed analysis at 5 months of age. qRT-PCR of whole kidneys showed reduction of *Pkd1* or *Ankmy2* in the respective conditional knockout animals at 5 months (**Supplemental Figure 3A**). At this age, *Pax8^rtTA^*; *TetO-Cre*; *Pkd1^f/f^* mice showed increased 2-kidney to body weight ratio and cystic index compared to controls (**Figure 3A-C, Supplemental 3B**). We noted a significant reduction in kidney to body weight ratio in *Pax8^rtTA^*; *TetO-Cre*; *Pkd1^f/f^*; *Ankmy2^f/f^* compared to *Pax8^rtTA^*; *TetO-Cre*; *Pkd1^f/f^* male animals. In addition, the cystic index was significantly reduced in the *Pax8^rtTA^*; *TetO-Cre*; *Pkd1^f/f^*; *Ankmy2^f/f^* compared to *Pax8^rtTA^*; *TetO-Cre*; *Pkd1^f/f^* mice (**Figure 3A-C**). *Pax8^rtTA^*; *TetO-Cre*; *Ankmy2^f/f^* kidneys remained unaffected. Immunostaining of kidney sections confirmed cysts in both LTL+ proximal tubules and AQP2+ collecting ducts in *Pax8^rtTA^*; *TetO-Cre*; *Pkd1^f/f^* that was reversed in *Pax8^rtTA^*; *TetO-Cre*; *Pkd1^f/f^*; *Ankmy2^f/f^* male mice (**Figure 3E-F, Supplemental Figure 3B**). Cyst sizes in *Pax8^rtTA^*; *TetO-Cre*; *Pkd1^f/f^* were impacted in both LTL+ and AQP2+ cysts compared to *Pax8^rtTA^*; *TetO-Cre*; *Pkd1^f/f^; Ankmy2^f/f^* but more so in LTL+ collecting duct cysts, which showed more differences between the two genotypes. The BUN values of *Pax8^rtTA^*; *TetO-Cre*; *Pkd1^f/f^*was significantly higher than *Pax8^rtTA^*; *TetO-Cre*; *Pkd1^f/f^*; *Ankmy2^f/f^* male mice, and the levels correlated with cyst severity (**Figure 3G**). However, kidney to body weight ratio, cystic index and cyst size of both LTL+ and AQP2+ cysts were not significantly different between *Pax8^rtTA^*; *TetO-Cre*; *Pkd1^f/f^*and *Pax8^rtTA^*; *TetO-Cre*; *Pkd1^f/f^*; *Ankmy2^f/f^* female mice (**Supplemental Figure 4A-D**), unlike in males. The lack of effect in female in contrast to male mice suggested sexual dimorphism in effects of ANKMY2 in the PKD model most likely from androgen receptor mediated control of sexually dimorphic gene expression in the proximal tubule.^50, 51^ Overall, lack of ANKMY2 suppressed adult onset cystogenesis in male mice.

**Figure 3.**
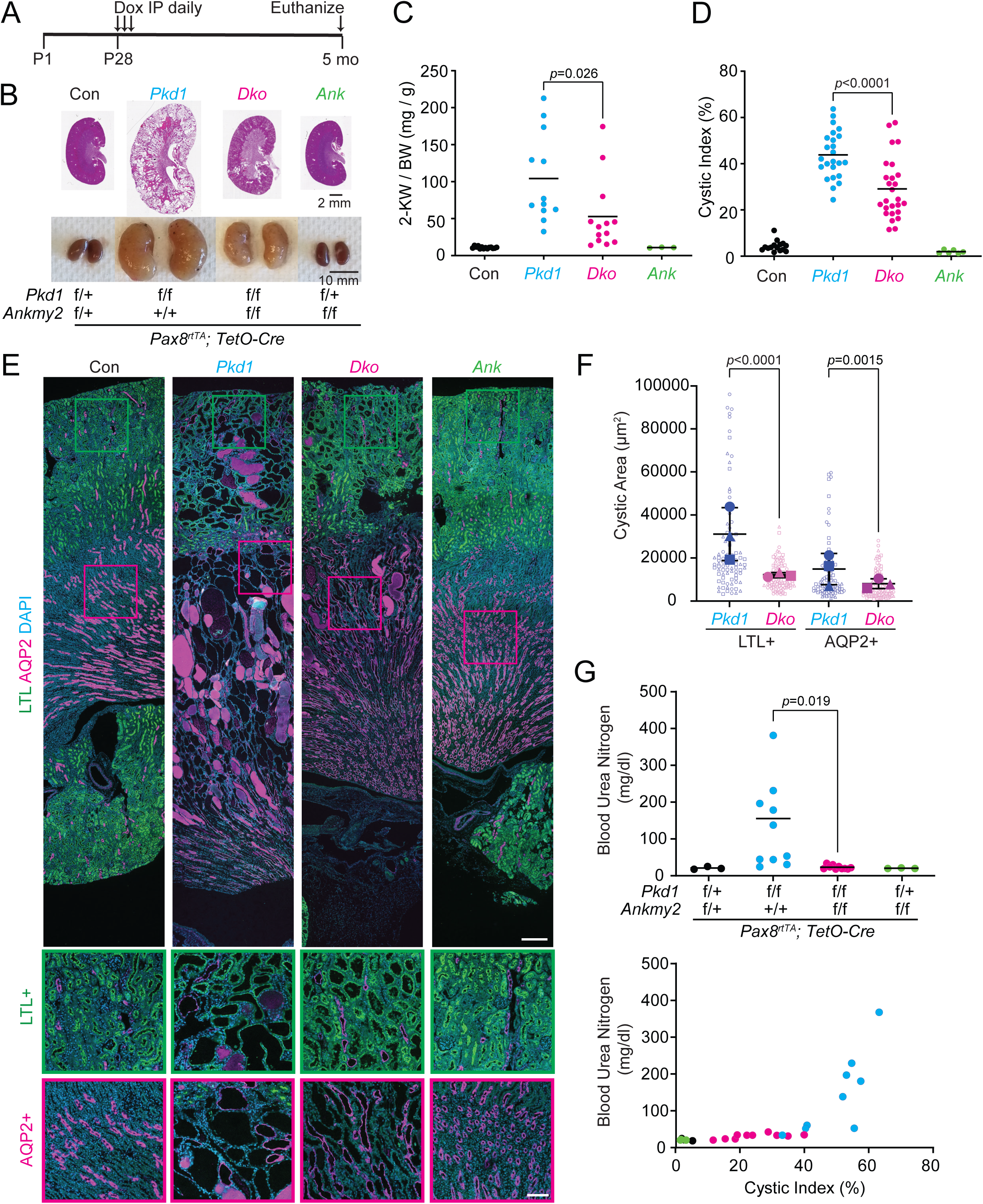
Cystogenesis in adult-onset PKD in male mice is suppressed from lack of ANKMY2. **(A)** Scheme for inducible conditional knockout in kidney nephron epithelia in an adult-onset model of PKD. **(B)** H&E and whole mount images of 5-month-old kidneys in control, *Pax8^rtTA^*; *TetO-Cre*; *Pkd1^f/f^*, *Pax8^rtTA^*; *TetO-Cre*; *Pkd1^f/f^; Ankmy2^f/f^*, *Pax8^rtTA^*; *TetO-Cre*; *Ankmy2^f/f^* male mice in C57BL/6J background. Images of multiple kidneys in male mice are shown in Figure S3. **(C)** 2-Kidney to body weight ratios of 5-month-old *Pax8^rtTA^*; *TetO-Cre*; *Pkd1^f/f^*male mice were significantly higher than *Pax8^rtTA^*; *TetO-Cre*; *Pkd1^f/f^; Ankmy2^f/^*. **(D)** Significantly increased cystic index in *Pax8^rtTA^*; *TetO-Cre*; *Pkd1^f/f^* male mice compared to *Pax8^rtTA^*; *TetO-Cre*; *Pkd1^f/f^; Ankmy2^f/f^*. **(E)** AQP2/LTL co-staining in kidney sections at 5 months shows multiple LTL positive cysts in cortical regions and corticomedullar junctions and AQP2 positive cysts in medullar, corticomedullar junctions and cortical regions in *Pax8^rtTA^*; *TetO-Cre*; *Pkd1^f/f^* animals compared to *Pax8^rtTA^*; *TetO-Cre*; *Pkd1^f/f^; Ankmy2^f/f^*. Scale: 300 µm. Magnified images of cortical and cortico-medullar junctions are shown below; LTL positive, green outlines; AQP2 positive, magenta outlines. Scale: 100 µm. **(F)** Cyst sizes in *Pax8^rtTA^*; *TetO-Cre*; *Pkd1^f/f^* were increased in both LTL+ and AQP2+ cysts compared to *Pax8^rtTA^*; *TetO-Cre*; *Pkd1^f/f^; Ankmy2^f/f^* but more so in LTL+ collecting duct cysts, which showed more differences between the two genotypes. Superplots of N=3 kidneys/genotype are shown with different shapes and averages are plotted with larger shapes. Cystic indices in *Pax8^rtTA^*; *TetO-Cre*; *Pkd1^f/f^* were 63, 47, and 40, whereas that in *Pax8^rtTA^*; *TetO-Cre*; *Pkd1^f/f^; Ankmy2^f/f^* were 22, 15, and 18. **(G)** The BUN values of *Pax8^rtTA^*; *TetO-Cre*; *Pkd1^f/f^* was significantly higher than *Pax8^rtTA^*; *TetO-Cre*; *Pkd1^f/f^*; *Ankmy2^f/f^* mice, and the levels correlated with cystic index across genotypes. See also Supplemental Figure 3 and 4.

### Increased proliferation in adult-onset PKD is suppressed from lack of ANKMY2

We next tested proliferation in the adult-onset PKD models. We immunostained kidney sections for Ki67 and quantified the percentage of cycling cells by counting Ki67+ epithelial cells co-stained with LTL (proximal tubule) or AQP2 (collecting duct). For this quantification, we tested kidneys from male mice with cystic indices close to the median values of respective genotypes. Ki67+ cyclic cells in proximal tubule and collecting duct epithelia were significantly increased in *Pax8^rtTA^*; *TetO-Cre*; *Pkd1^f/f^* compared to *Pax8^rtTA^*; *TetO-Cre*; *Pkd1^f/f^*; *Ankmy2^f/f^*(**Figure 4A-C**). Cyst formation in conditional knockouts of *Pkd1/2* is associated with activation of components of the extracellular regulated kinase (ERK) pathway.^10, 52^ The amounts of phosphorylated and total ERK in whole kidney lysates increased in *Pax8^rtTA^*; *TetO-Cre*; *Pkd1^f/f^* cystic kidneys compared to *Pax8^rtTA^*; *TetO-Cre*; *Pkd1^f/f^*; *Ankmy2^f/f^* kidneys (**Figure 4D**). Finally, as an orthogonal measure of cyst severity, we checked the transcript levels of Kidney Injury Molecule 1 (KIM-1). KIM1 is a transmembrane glycoprotein that is expressed mostly in proximal tubular cells and is elevated early in tubulointerstitial injury in both rodent models ^53, 54^ and in humans.^55^ Levels of *Kim1* transcripts significantly increased in *Pax8^rtTA^*; *TetO-Cre*; *Pkd1^f/f^* cystic kidneys compared to *Pax8^rtTA^*; *TetO-Cre*; *Pkd1^f/f^*; *Ankmy2^f/f^* kidneys and remained unchanged in *Pax8^rtTA^*; *TetO-Cre*; *Ankmy2^f/f^* kidneys compared to controls (**Figure 4E**). These results suggest that cyst burden is suppressed in adult-onset PKD from lack of ANKMY2 from decreased proliferation, reduced ERK activation, and exhibits reduced expression of tubulointerstitial injury marker KIM-1.

**Figure 4.**
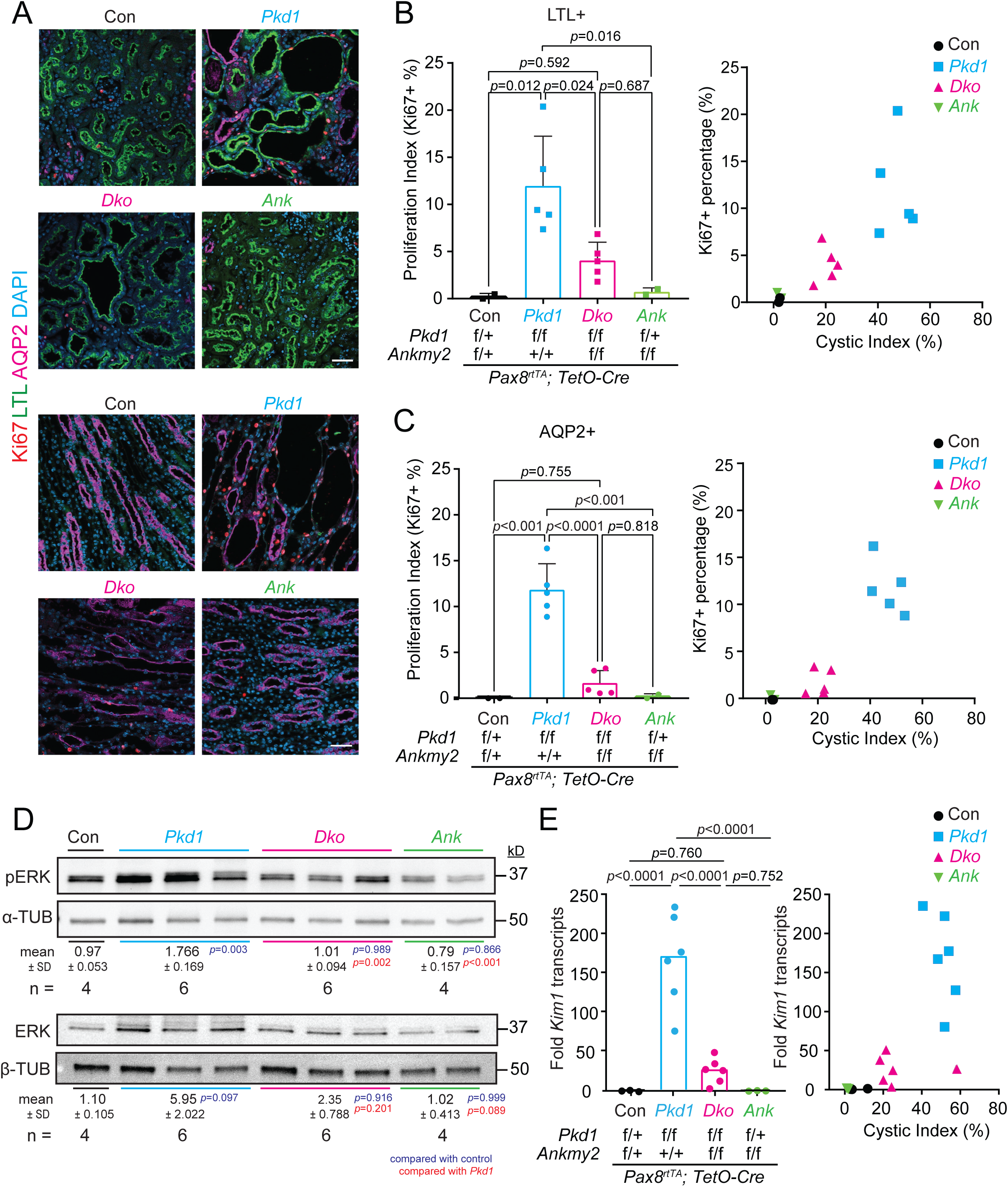
Increased proliferation in adult-onset PKD in male mice is suppressed from lack of ANKMY2. **(A-C)** Increased proliferation in cystic kidneys in 5-month-old *Pax8^rtTA^*; *TetO-Cre*; *Pkd1^f/f^* male mice compared to *Pax8^rtTA^*; *TetO-Cre*; *Pkd1^f/f^; Ankmy2^f/f^*. Kidney sections were immunostained for Ki67, AQP2 and LTL and counterstained with DAPI. Ki67 + cells were quantified. Regression analysis with respect to cystic index is shown. **(D)** Immunoblotting for pERK and ERK, of (i) control, *Pax8^rtTA^*; (ii) *TetO-Cre*; *Pkd1^f/f^*, *Pax8^rtTA^*; *TetO-Cre*; (iii) *Pkd1^f/f^; Ankmy2^f/f^* and *Pax8^rtTA^*; (iv) *TetO-Cre*; *Ankmy2^f/f^* kidneys from male mice at 5 months. Loading controls (Tubulin) for each phosphoprotein is shown. Levels of individual phosphoprotein or total protein, each normalized to tubulin separately, is shown. Control, N=4; *Pax8^rtTA^*; *TetO-Cre*; *Pkd1^f/f^* N=6; *Pkd1^f/f^; Ankmy2^f/f^* and *Pax8^rtTA^* N=6; and *TetO-Cre*; *Ankmy2^f/f^* N=4. **(E)** Transcript levels of *Kim1* in whole kidneys from *Pax8^rtTA^*; *TetO-Cre*; *Pkd1^f/f^*are significantly higher than *Pax8^rtTA^*; *TetO-Cre*; *Pkd1^f/f^; Ankmy2^f/f^*, and the levels correlated with cystic index across genotypes.

### Ciliary pools of ADCY5 and ADCY6 are reduced in *Ankmy2* knockout kidney epithelial cells

We next determined the mechanism by which ANKMY2 was regulating PKD cystogenesis. We previously showed that lack of ANKMY2 prevents localization of stably expressed ADCY3/5/6 in cilia of mouse fibroblasts and in neuroepithelial cells without grossly affecting cilia architecture.^46^ Of the nine membrane adenylyl cyclases, ADCY6 is the predominant adenylyl cyclase detected throughout the nephron, whereas ADCY5 is the second most prevalent adenylyl cyclase detected in the connecting tubule and cortical collecting duct using proteomic studies of the nephron segments.^56^ As ADCY5 and ADCY6 are expressed in kidney epithelia, we stably expressed LAP-tagged versions in mouse IMCD3-FlpIn cells. Both ADCY5^LAP^ and ^LAP^ADCY6 localized to IMCD3-FlpIn cilia, although ^LAP^ADCY6 positive ciliary percentage was less than in the previously reported NIH 3T3 cells ^46^ (**Figure 5A-D**). In addition, both these adenylyl cyclases were also localized to the secretory pathway and plasma membranes (**Supplemental Figure 5A-B**). We next generated *Ankmy2* knockouts in the ADCY5^LAP^ and ^LAP^ADCY6 IMCD3 cells that was confirmed from the complete loss of ANKMY2 using immunoblotting for ANKMY2 (**Figure 5G, Supplemental Figure 5C**). Interestingly, we noted a significant decrease of ADCY5^LAP^ and ^LAP^ADCY6 ciliary and plasma membrane levels in *Ankmy2* ko cells despite persistent extraciliary localization in the secretory pathway (**Figure 5A-D, Supplemental Figure 5A-B**). Gross ciliary morphology or localization of other TULP3 cargoes such as GPR161 or other non-TULP3 trafficked cilia localized proteins such as the hedgehog pathway transducer—SMO remained unaffected in *Ankmy2* ko IMCD3 cells.^46^ We previously showed that complex glycosylation of adenylyl cyclases were affected in *Ankmy2* ko fibroblasts.^46^ Both ADCY5 and ADCY6 were in complex glycosylated forms in IMCD3 cells, as determined by EndoH and PNGase treatments (**Figure 5E-F**). The levels of complex glycosylated forms of ADCY5/6 were reduced in *Ankmy2* ko IMCD3-FlpIn cells despite equivalent expression but were restored upon stable reexpression of ^HA^ANKMY2 (**Figure 5G**). Such reexpression also restored plasma membrane levels of ADCY5^LAP^ and ^LAP^ADCY6 (**Supplemental Figure 5A-B**). Although reexpression did not rescue ciliary levels of ADCY5^LAP^ and ^LAP^ADCY6 in IMCD3 cells, we previously showed that ciliary levels were partially restored in mouse fibroblasts.^46^ Such lack of rescue of ciliary levels is likely from extensive plasma membrane localization in IMCD3-FlpIn vs 3T3-FlpIn cells. Thus, lack of ANKMY2 prevents ciliary localization of ADCY5^LAP^ and ^LAP^ADCY6 while still retaining cellular levels. In contrast, in the *Adcy5/6* knockouts, total cellular levels of respective adenylyl cyclases would be absent. Therefore, conditional knockouts of *Ankmy2* prevented ciliary compartmentalization of multiple adenylyl cyclases in the kidney epithelia.

**Figure 5.**
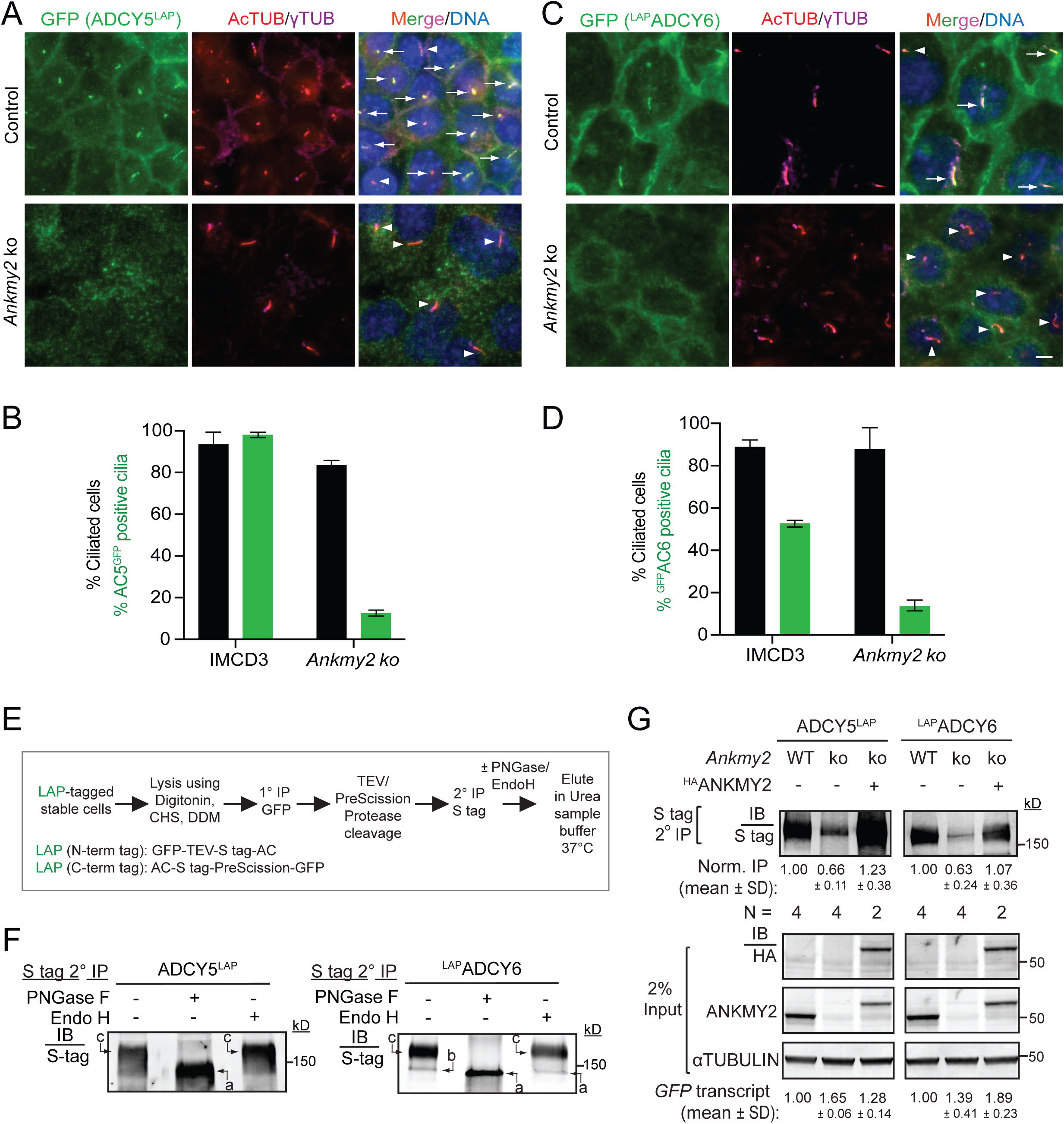
Ciliary pools of ADCY5 and ADCY6 are reduced in *Ankmy2* knockout kidney epithelial cells. **(A-D)** ADCY5^LAP^ and ^LAP^ADCY6 were localized to cilia in stably expressing IMCD3 FlpIn cell line and lacked them in cilia upon CRISPR-based *Ankmy2* ko, as detected upon performing immunostaining with antibodies against GFP and acetylated tubulin. Scale: 5 µm. Quantification in (B, D) shown as mean ± SD. Total 225-500 cells counted/cell line from 2-3 separate images. Arrows and arrowheads show GFP+ and GFP-cilia, respectively. **(E-F)** Immunoblots showing glycosylation state of stably expressed LAP-tagged Adenylyl cyclases in IMCD3 FlpIn cells after tandem affinity purification followed by Endo H/PNGase treatment (flowchart in E). Form “c”, complex glycosylated; Form “b”, core glycosylated; Form “a”, non-glycosylated. Abbreviations: DDM, n-Dodecyl-β-D-Maltoside; CHS, Cholesteryl hemisuccinate. **(G)** Immunoblots showing glycosylated states of stably expressed LAP-tagged AC5 and AC6 present in control and *Ankmy2* ko ± ^HA^ANKMY2 IMCD3 FlpIn cells as shown in (E). Immunoblots for ANKMY2 and ^HA^ANKMY2 in lysates along with α-tubulin are shown below. Data showing ratios of complex vs core glycosylated forms (c/b) from N=2-4 experiments, expressed as mean ± SD. P values with respect to control cells. See also Supplemental Figure 5.

### Ciliary length increase from PC1 loss in kidney epithelia is suppressed from ANKMY2 loss

Optogenetic activation of a cilia targeted bacterial photoactivable adenylyl cyclase (bPAC) elevates ciliary cAMP levels and results in a corresponding increase in ciliary length after activation for an hour.^57, 58^ As ANKMY2 regulated ciliary localization of adenylyl cyclases to cilia, we next tested cilia changes in the adult-onset PKD models. We reasoned that changes in cilia length in an ANKMY2-dependent manner accompanying cystogenesis would confirm involvement of ciliary adenylyl cyclase signaling. We co-immunostained kidney sections with cilia marker acetylated tubulin in epithelial LTL- or AQP2-stained cells to quantify ciliary length in proximal tubule and collecting duct epithelia, respectively (**Figure 6A-B**) at 5 months (mice were treated like in Figure 3A). For this quantification, we chose kidneys from male and female mice of respective genotypes with cystic indices across the spectrum (**Figure 6C**). Cilia lengths were lower in the proximal tubules compared to the collecting ducts, as reported before.^59^ We found that kidney epithelia of both male and female *Pax8^rtTA^*; *TetO-Cre*; *Pkd1^f/f^* mice showed long cilia compared to controls in both proximal tubule and collecting duct regions (**Figure 6A-C, Supplemental Figure 6A**). Such increase was apparent in both male and female animals (**Figure 6C, Supplemental Figure 6A**). Interestingly, *Pax8^rtTA^*; *TetO-Cre*; *Pkd1^f/f^*; *Ankmy2^f/f^* showed significantly reduced ciliary length compared to *Pax8^rtTA^*; *TetO-Cre*; *Pkd1^f/f^* male and female animals in both proximal tubule and collecting duct regions (**Figure 6C**). Furthermore, the cilia length in the *Pax8^rtTA^*; *TetO-Cre*; *Pkd1^f/f^*; *Ankmy2^f/f^* male and female animals remained unaffected by cyst severity, whereas the cilia length in *Pax8^rtTA^*; *TetO-Cre*; *Pkd1^f/f^* male and female animals scaled with cystic index increase (**Figure 6C**). In contrast, total cAMP levels were reflective of cyst severity irrespective of genotype or sex, suggesting that increased cellular cAMP levels in the *Pax8^rtTA^*; *TetO-Cre*; *Pkd1^f/f^*; *Ankmy2^f/f^* were insufficient to induce ciliary length increase (**Figure 6D**). Therefore, lack of ANKMY2 blocks ciliary length increase in *Pax8^rtTA^*; *TetO-Cre*; *Pkd1^f/f^* animals, suggesting that ciliary adenylyl cyclase signaling mediates increased cilia length in ADPKD mice models.

**Figure 6.**
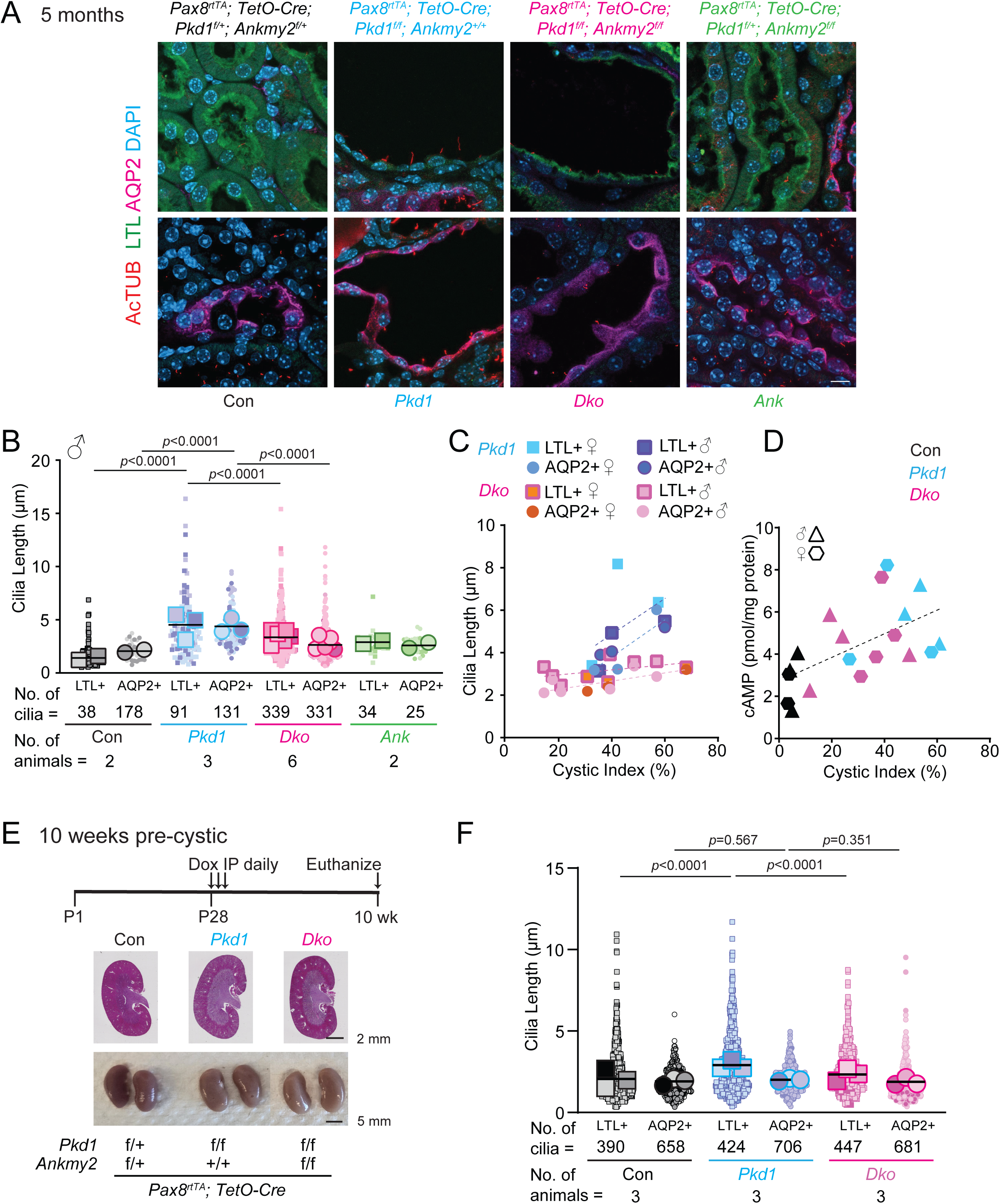
Ciliary length increase from PC1 loss in adult kidney epithelia is suppressed from ANKMY2 loss. **(A)** Kidney sections of 5-month-old mice (as in Figure 3a) were immunostained for acetylated tubulin, AQP2 and LTL and counterstained with DAPI. Scale 10 μm. **(B)** Cilia lengths were quantified and super plots of N=2-6 male 5-month-old mice shown for each genotype. Lengths from each kidney is shown with different shapes and averages are plotted with larger shapes. Data for female mice is shown in Supplemental Figure 6A. **(C)** Mean cilia lengths with respect to cystic indices shown for 5-month-old mice. Cilia length in *Pax8^rtTA^*; *TetO-Cre*; *Pkd1^f/f^* animals scale with cystic indices but remains unchanged irrespective of cystic indices in *Pax8^rtTA^*; *TetO-Cre*; *Pkd1^f/f^; Ankmy2^f/f^*. Both males and females are shown. **(D)** Total cAMP levels with respect to cystic indices shown for respective genotypes for 5-month-old mice and the levels correlated with cystic index across genotypes irrespective of sex. **(E)** Scheme for inducible conditional knockout in kidney nephron epithelia in an adult-onset model of PKD to detect “pre-cystic” tubules. H&E and whole mount images of 10-week-old kidneys in control, *Pax8^rtTA^*; *TetO-Cre*; *Pkd1^f/f^*and *Pax8^rtTA^*; *TetO-Cre*; *Pkd1^f/f^; Ankmy2^f/f^* male mice in C57BL/6J background. Images of multiple kidneys, 2-kidney to body weight ratios, and cystic indices in male mice are shown in Figure S6B. **(F)** Cilia lengths were quantified and super plots of N=3 male 10-week-old animals shown for each genotype. Lengths from each kidney is shown with different shapes and averages are plotted with larger shapes. Immunofluorescence images are shown in Supplemental Figure 6B. See also Supplemental Figure 6.

### Ciliary length increase from PC1 loss in kidney epithelia precedes cystogenesis

Cilia length increases in *Pax8^rtTA^*; *TetO-Cre*; *Pkd1^f/f^* mice could be a cause or consequence of cystogenesis. Cilia length changes in *Pax8^rtTA^*; *TetO-Cre*; *Pkd1^f/f^*mice preceding cyst progression would argue for ciliary involvement preceding cystogenesis. Therefore, we checked kidneys at 10 weeks following Doxycycline injections at P28-30 and measured cilia lengths in proximal tubules and collecting ducts (**Figure 6E**). The kidney tubules in *Pax8^rtTA^*; *TetO-Cre*; *Pkd1^f/f^*mice just started to show dilations at this stage as reported earlier,^60^ but overt cysts were rare (**Supplemental Figure 6B**), a condition we call “pre-cystic”. At this stage the 2-kidney/body weight ratios and kidney cystic indices did not show significant differences between *Pax8^rtTA^*; *TetO-Cre*; *Pkd1^f/f^* and *Pax8^rtTA^*; *TetO-Cre*; *Pkd1^f/f^*; *Ankmy2^f/f^* males (**Figure 6E, Supplemental Figure 6B**). However, we found that the cilia lengths in the proximal tubule epithelial cells of *Pax8^rtTA^*; *TetO-Cre*; *Pkd1^f/f^* males were significantly higher than control or *Pax8^rtTA^*; *TetO-Cre*; *Pkd1^f/f^*; *Ankmy2^f/f^* males (**Figure 6F**). Although the cilia lengths in proximal tubule epithelia of *Pax8^rtTA^*; *TetO-Cre*; *Pkd1^f/f^* males were not as high as at 5 months (**Figure 6B**), the process had initiated by 10 weeks. We did not capture changes in cilia lengths in collecting duct epithelial cilia at this early stage (**Figure 6F**). Nonetheless, the initiation of cilia length increases at the pre-cystic stage in proximal tubule epithelia of *Pax8^rtTA^*; *TetO-Cre*; *Pkd1^f/f^* males suggest ciliary length changes preceding cystogenesis. Furthermore, the lack of ciliary length increases in absence of *Ankmy2* suggest permissive changes in cilia from ciliary adenylyl cyclase targeting that ultimately dictate the massive ciliary length increase in *Pax8^rtTA^*; *TetO-Cre*; *Pkd1^f/f^* males. These results suggest cilia involvement preceding cystogenesis.

## Discussion

Our results show that lack of ANKMY2 can suppress both embryonic-onset or adult-onset cystogenesis. However, lack of ANKMY2 only works at an early postnatal window and such suppression is overridden later, presumably from downstream cellular cAMP signaling that is not exquisitely ANKMY2 regulated. In fact, pCREB levels in collecting duct epithelia in later postnatal kidneys at P15 are similar between *Hoxb7-Cr*e; *Pkd1^f/f^* and *Hoxb7-Cre*; *Pkd1^f/f^*; *Ankmy2 ^f/f^* animals. Thus, the lack of ciliary adenylyl cyclase signaling from ANKMY2 loss can only suppress early cystogenesis in collecting ducts but could be ineffective at later stages when cellular cAMP levels are active during cyst progression. Loss of *TULP3* or IFT-A subunit *Thm1* exacerbates PKD from concomitant deletion of *Pkd1/2* at the juvenile stage.^12, 13^ Part of this exacerbation is likely from lack of TULP3 cargo ARL13B from cilia at this stage.^15, 16^ In contrast, our results with *Ankmy2* conditional knockout suggest that loss of adenylyl cyclases from cilia prevents early embryonic onset cystogenesis and improves life expectancy but is unable to revert final cyst progression. Loss of ANKMY2 also prevents cystogenesis in adult-onset ADPKD models in male mice but not in female mice. Cyst reduction was impacted in both proximal tubules and collecting ducts, but more so in proximal tubules. Human ADPKD and rodent models display sexual dimorphism in renal disease progression secondary to differences in the levels of gonadal hormones.^61, 62^ Ancillary factors that prevent cyst regulation by ciliary adenylyl cyclase signaling in female kidneys could encompass androgen receptor mediated control of sexually dimorphic gene expression in the proximal tubule.^50, 51^

Mechanistically, lack of ANKMY2 prevented localization of multiple adenylyl cyclases to cilia while still retaining mature glycoslylated forms. We find that cilia lengths are drastically increased in adult-onset *Pkd1* conditional ko cystic kidneys (*Pkd1*) in an ANKMY2-dependent manner. We suggest de-repression of an ANKMY2-dependent ciliary cAMP pathway preceding cystogenesis in adult-onset *Pax8^rtTA^*; *TetO-Cre*; *Pkd1^f/f^* mice (**Figure 7**) for the following reasons. First, cilia length increase in adult ADPKD models was suppressed from ANKMY2 loss, irrespective of cyst load or sexual dimorphism in cyst suppression. Second, cilia length increases initiated at the pre-cystic stage in proximal tubule epithelia of *Pax8^rtTA^*; *TetO-Cre*; *Pkd1^f/f^* males suggesting ciliary length changes preceding cystogenesis. Third, cilia length in adult ADPKD models remained unchanged from loss of ANKMY2 despite high cystic load suggesting that cytoplasmic cAMP was insufficient to trigger cilia length increase. Rather, the lack of ciliary length increases in absence of *Ankmy2* suggest permissive changes from ciliary adenylyl cyclase targeting during the pre-cystic stage that ultimately dictate the massive ciliary length increase in *Pax8^rtTA^*; *TetO-Cre*; *Pkd1^f/f^*. Thus, the overtly long cilia in *Pax8^rtTA^*; *TetO-Cre*; *Pkd1^f/f^* mice at 5 months are likely from initiation of ciliary length increase starting from pre-cystic conditions and high ciliary cAMP diffusing from cytoplasm at later cystic stages. Overall, our results suggest both cilia-dependent and cilia-independent mechanisms for cAMP regulation during renal cyst initiation and progression.

**Figure 7.**
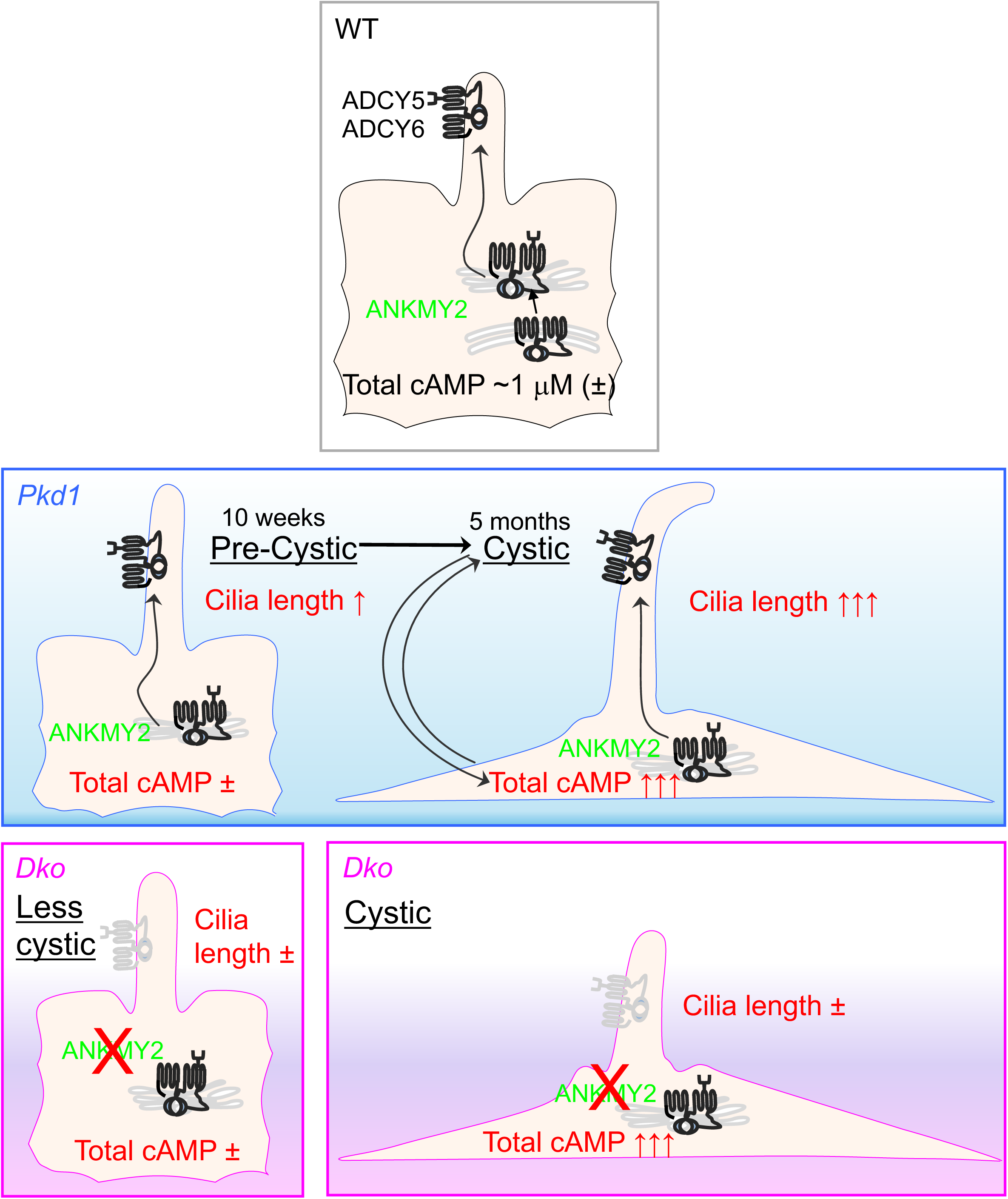
Model showing de-repression of an ANKMY2-dependent ciliary cAMP pathway preceding cystogenesis in adult-onset PKD. In wild-type, ANKMY2 functions in maturation and localization of adenylyl cyclases to kidney epithelial cilia. Optogenetic activation of a photoactivable adenylyl cyclase inside cilia increases cilia length, suggesting that cAMP levels in cilia determine cilia length.^57, 58^ We find that cilia lengths are drastically increased in adult-onset *Pax8^rtTA^*; *TetO-Cre*; *Pkd1^f/f^* cystic kidneys (*Pkd1*) at 5 months. At 10 weeks in pre-cystic kidneys, we note initiation of cilia length increase in these animals, suggesting that ciliary changes precede cyst formation. Thus, the overtly long cilia in *Pkd1* mice at 5 months are likely from initiation of ciliary length increase starting from pre-cystic conditions and high ciliary cAMP diffusing from cytoplasm at later cystic stages. In adult-onset *Pax8^rtTA^*; *TetO-Cre*; *Pkd1^f/f^*; *Ankmy2^f/f^* (*Dko*) kidneys the cilia lengths are never increased, even in the few male mice with bigger cysts, or in female mice with overt cysts, suggesting that high cellular cAMP levels are unable to increase cilia length in the absence of ANKMY2. These results suggest de-repression of an ANKMY2-dependent ciliary cAMP pathway preceding cystogenesis in *Pkd1* mice. Furthermore, permissive changes from ciliary adenylyl cyclase targeting during pre-cystic stage ultimately dictate the massive ciliary length increase in *Pkd1* mice.

Ciliary length is exquisitely controlled in diverse organisms and tissues.^63, 64^ Cilia lengths are lower in the proximal tubules compared to the collecting ducts, and reduce with age.^59^ Our results show that one critical index of ciliary regulation involves maintenance of ciliary length by polycystin signaling during adult-onset ADPKD initiation and progression in an ANKMY2-dependent manner. As loss of ANKMY2 results in reduced AC5/6 levels in cilia, taken together, our results implicate ciliary adenylyl cyclase signaling in increase in ciliary length from *Pkd1* deletion. Loss of PC1/2 has been shown to increase cilia length in ADPKD mice models^12, 65, 66^ and in human patients,^67^ although there are conflicting results in cell lines showing both increased^68^ or decreased cilia lengths.^69^ Increased *Pkd1* dosage also induces increased cilia length in both precystic and cystic tubules.^70^ Thus, PC1 dosage is associated with variable changes in cilia length in different model systems. Importantly, optogenetic activation of bPAC elevates ciliary cAMP levels and results in a corresponding increase in ciliary length after activation for an hour.^57, 58^ Elevated acute cellular cAMP signaling has been reported to increase ciliary length in cultured mammalian kidney epithelial cells.^71^ Importantly, while total cAMP levels were reflective of cyst severity irrespective of genotype or sex, increased cellular cAMP levels were unable to mediate ciliary length increase in absence of ANKMY2. The cAMP-mediated ciliary length control in cultured mammalian kidney epithelial cells is also regulated by flow^71^ mimicking the nephron lumen,^59^ such that increased ciliary length from forskolin treatment is prevented in the presence of flow.^71^ It is possible that pathological increase in cilia length upon *Pkd1* deletion prevents flow-mediated ciliary length control, thereby establishing a forward feedback effect in reinforcing increased cilia length.

How does ciliary cAMP signaling regulate kidney tubule homeostasis? Ciliary and cytoplasmic cAMP levels are differentially interpreted in vertebrate cells in hedgehog signaling^33^ and in determining tissue-specific phenotypes.^72, 73^ The only experiment till date that has tested ciliary cAMP pools in cystogenesis was by chronic optogenetic stimulation of cilia targeted bacterial photoactivable adenylyl cyclase (bPAC).^57, 58^ A bPAC specifically targeted to cilia results in cystic changes in 3D cultured cells, but not upon being only targeted to cytoplasm.^57, 58^ Increased ciliary length possibly regulates ciliary and periciliary effectors in downstream regulation of pathways, most likely pivotal in early response of transcriptional and post-transcriptional targets that initiate cystogenesis. There is precedence to downstream transcriptional regulation as for phase regulation of the core transcriptional circuit in the circadian clock by diurnal fluctuations of ciliary lengths in hypothalamic suprachiasmatic nuclei.^74^ Similarly, post-transcriptional regulation of targets by mechanical transduction in crown cilia in the embryonic node is pivotal in left right asymmetry.^75, 76^ The local diffusion of cAMP from cilia^76^ and local buffering from phosphodiesterases^37, 77^ could result in periciliary surges in regulating effectors. The ciliary cAMP-mediated early transcriptional and post-transcriptional targets during ADPKD cystogenesis are currently unknown and will be the focus of future studies. Nonetheless, targeting early initiation/progression of cyst pathogenesis along with existing drugs that target cellular cAMP levels^31^ can fundamentally change the treatment landscape of ADPKD. Finally, our results provide a framework for identifying and testing ciliary effectors and early subcellular events that initiate cystogenesis in the future.

## Materials and Methods

### Mouse strains and genotyping

All mice were housed at the Animal Resource Center of the University of Texas Southwestern (UTSW) Medical Center. Mice were housed in standard cages that contained three to five mice per cage, with water and standard diet *ad libitum* and a 12 h light/dark cycle. Both male and female mice were analyzed in all experiments. *HoxB7-Cre* mice were obtained from O’Brien Kidney Research Core of UT Southwestern. *Pkd1^f/f^* allele has been described before^52^ and were backcrossed to C57BL/6J background. *Pax8^rtTA^*; *TetO-Cre*^10^ combined with *Pkd1^f/f^* allele^78^ that were back crossed to C57BL/6J were obtained from PKD RRC, University of Maryland, School of Medicine. ES cells targeting *Ankmy2* (NM_146033.3, MGI: 2144755; third exon flanked by LoxP sites) that was generated by homologous recombination in mouse ES cells of the C57BL/6 strain were from EUCOMM (Clone # HEPD0679-6-C03)^46^. Mice were housed in standard cages that contained three to five mice per cage, with water and standard diet *ad libitum* and a 12 h light/dark cycle. All protocols were approved by the UTSW Institutional Animal Care and Use Committee. All the animals in the study were handled according to protocols approved by the UT Southwestern Institutional Animal Care and Use Committee, and the mouse colonies were maintained in a barrier facility at UT Southwestern, in agreement with the State of Texas legal and ethical standards of animal care.

### Mouse ES cells

ES cells targeting *Ankmy2* (NM_146033.3, MGI: 2144755; third exon flanked by LoxP sites) that was generated by homologous recombination in mouse ES cells of the C57BL/6 strain were from EUCOMM (Clone # HEPD0679-6-C03)^46^. ES cells were grown on SNL feeders with media containing 20% Serum, 6 mM L-glutamine, 1X Penicillin/Streptomycin, 1 mM β-mercaptoethanol, 1 mM Non-essential Amino Acids, 1X Nucleosides, 10 mg/L Sodium Pyruvate, ESGRO supplement 66 μl/L and incubated at 37 ^0^C in 5% CO_2_ (Dr. Robert Hammer lab, UT Southwestern Medical Center, Dallas). The ES cells were injected into host embryos of the C57BL/6 albino strain by the transgenic core (Dr. Robert Hammer lab, UT Southwestern Medical Center, Dallas

### Mouse genotyping

To genotype *Ankmy2* mice, following primers were used. For floxed allele with or without deletion: 3F (5’-CTG TCT CCA TAT TCA CAC ATT GAA TAG C-3’), 4R (5’-GCT GCA TGC ATC AAA GGA GTC ATT CC-3’) and 2R (5’-TGA ACT GAT GGC GAG CTC AGA CC-3’) gave 508 bp for wild type, 732 bp for floxed allele, and 289 bp for deleted floxed allele (cko). *Cre* allele was genotyped with Cre-F (5’-AAT GCT GTC ACT TGG TCG TGG C-3’) and Cre-R (5’-GAA AAT GCT TCT GTC CGT TTG C-3’) primers (100 bp amplicon). For the *Pkd1 ^tm2Som^* ^52^ allele, forward (5’-CCG CTG TGT CTC AGT GTC TG-3’) and reverse (5’-CAA GAG GGC TTT TCT TGC TG-3’) gave 400 bp for floxed allele and 200 bp for wild type. For the *Pkd1^tm2Ggg^*allele^78^, F4 forward (5’-CCT GCC TTG CTC TAC TTT CC-3’) and R5 back (5’-AGG GCT TTT CTT GCT GGT CT-3’) gave 250 bp for floxed allele and 180 bp for wild type. For *Pax8^rtTA^*, oIMR7385 (5’-CCA TGT CTA GAC TGG ACA AGA-3’) and oIMR7386 (5’-CTC CAG GCC ACA TAT GAT TAG-3’) gave 595 bp for *Pax8^rtTA^* allele.

### Cell culture and generation of stable cell lines

IMCD3 Flp-In, Phoenix A (PhA, Indiana University National Gene Vector Biorepository), and 293FT cells were cultured in DMEM high glucose (Sigma-Aldrich; supplemented with 10% cosmic serum, 0.05 mg/ml penicillin, 0.05 mg/ml streptomycin, and 4.5 mM glutamine). They have tested negative for Mycoplasma using the Mycoplasma PCR Detection Kit (Genlantis). IMCD3 Flp-In, Phoenix A (PhA, Indiana University National Gene Vector Biorepository), and 293FT cells were cultured in DMEM high glucose (Sigma-Aldrich; supplemented with 10% cosmic serum, 0.05 mg/ml penicillin, 0.05 mg/ml streptomycin, and 4.5 mM glutamine). They have tested negative for Mycoplasma using the Mycoplasma PCR Detection Kit (Genlantis). Transfection of plasmids was done with Polyfect (QIAGEN) or Polyethylenimine (PEI) max. Stable cell lines were generated by retroviral infection or transfection. In many cases, stable lines were flow sorted and further selected for GFP. IMCD3 FlpIn cells stably expressing LAP tag or LAP tagged Adenylyl cyclases were generated by retroviral infection, antibiotic selection, and flow sorting. CRISPR/Cas9 knockout or knockdown lines for *Ankmy2* were generated in IMCD3 Flp-In cells by using guide RNA targeting sequences AAGGAACTGCTGGAAGTGAT (Exon 1) or GAATGTTCATGTCAACTGCT (Exon 2). Clonal lines were tested for knockout or knockdown by Sanger sequencing and immunoblotting for Ankmy2. ORF clones were as follows: ADCY5 (gift of Ron Taussig, UT Southwestern), ADCY6 (Life Technologies IOH40476), ANKMY2 (Origene RG206770; NM_020319 in PCMV6-AC-GFP vector from Origene). Gateway pENTR clones were generated by PCR cloning and BP reaction as necessary for N- or C-terminal tagging. Gatewaytized pBABE-LAP-N terminus and pBABE-LAP-C terminus plasmids were generated from LAP1 and LAP5 vectors (Addgene) and pBABE puro. We cloned 3×HA-ANKMY2 into pQXIN (Clontech), which was used for retroviral infection in knockout lines expressing LAP-tagged Adenylyl cyclases. Antibiotic selection was used to generate rescue lines stably expressing 3×HA-ANKMY2.

### Reverse transcription and quantitative PCR

RNA was extracted from using GenElute mammalian total RNA purification kit (RTN350; Sigma). qRT-PCR was performed with Kicqstart One-Step Probe RT-qPCR ReadyMix (KCQS07; Sigma). Inventoried TaqMan probes for qRT-PCR from Applied Biosystems were used. Reactions were run in CFX96 Real time System (Bio Rad).

### Tissue processing, antibodies, immunostaining and microscopy

Mice were perfused with PBS, and the kidneys were dissected and fixed in 4% paraformaldehyde overnight at 4°C and processed for cryosection or paraffin embedding and sectioning. For cryosection, the kidneys were incubated in 30% sucrose for 1-2 days at 4°C. Kidneys were mounted with OCT compound and cut into 15 µm frozen sections. For paraffin section, kidneys were processed over a 12-hour period using a Thermo-Fisher Excelsior Automated Tissue Processor (A82300001; ThermoFisher Scientific), which dehydrated the kidneys through 6 ethanol concentrations, from 50% ethanol to 100% ethanol, cleared through 3 changes of xylene, and infiltrated with wax through 3 Paraplast Plus paraffin baths (39602004; Leica). Samples were embedded in Paraplast Plus using paraffin-filled stainless steel base molds and a Thermo-Shandon Histocenter 2 Embedding Workstation (6400012D; ThermoFisher Scientific). The kidneys were then cut in 5 μm thick sections, deparaffined and treated with microwave in Antigen Retrieval citra solution (HK086-9K; BioGenex. Fremont, CA) for 10 min. For frozen sections, the sections were incubated in PBS for 15 min to dissolve away the OCT. Sections were then blocked in blocking buffer (1% normal donkey serum [Jackson immunoResearch, West Grove, PA], in PBS) for 1 hour at room temperature. Sections were incubated with primary antibodies against the following antigens; overnight at room temperature or 4C: Acetylated tubulin (T6793; Sigma mouse IgG2b, 1:500), AQP2 (SC515770, Santa Cruz Biotechnology, mouse IgG1, 1:500), Ki67 (ab16667, Abcam, 1:500), pCREB (9198S; Cell Signaling, 1:500). After three PBS washes, the sections were incubated in secondary antibodies (Alexa Fluor 488-, 555-, 594-, 647-conjugated secondary antibodies, or anti-mouse IgG isotype-specific secondary antibodies; 1:500; Life Technologies, Carlsbad, CA or Jackson ImmunoResearch) or cell surface markers, Fluorescein labeled Lotus tetragolonobus lectin (LTL; 1:200, FL 1321-2 Vector laboratories) for 1 hour at room temperature. Cell nuclei were stained with DAPI (Sigma) or Hoechst 33342 (Life technologies). Slides were mounted with Fluoromount-G (0100-01; Southern Biotech) and images were acquired with a Zeiss AxioImager.Z1 microscope or a Zeiss LSM980 confocal microscope. For hematoxylin and eosin staining, paraffin sections were stained by hematoxylin and eosin (Hematoxylin 560; 3801575; Leica and Alcoholic Eosin Y 515; 3801615; Leica) using a Sakura DRS-601 x-y-z robotic-stainer (DRS-601; Sakura-FineTek, Torrance, CA). Slides were dehydrated and mounted with Permount (SP15-100; ThermoFisher Scientific). For immunofluorescence experiments in cell lines, cells were cultured on coverslips until confluent and starved for indicated periods before fixation. Cells were fixed with 4% PFA. After blocking with 5% normal donkey serum, the cells were incubated with primary antibody solutions for 1 h at room temperature followed by treatment with secondary antibodies for 30 min along with DAPI. Primary antibodies used were against the following antigens: GFP (Abcam ab13970), Acetylated tubulin (T6793 Sigma mouse IgG2b, 1:500), γ-tubulin (sc-17787; Santacruz Biotech mouse IgG2a, 1:500), β-catenin (610154; BD Biosciences, 1:500), HA (clone 3F10; Roche, 1:500). Coverslips were mounted with Fluoromount-G and images were acquired with a Zeiss AxioImager.Z1 microscope.

### Cystic index quantification

Cystic index was quantified in paraffin embedded mid-sagittal sections of whole kidneys. HE stained kidney sections were photographed by the PrimeHisto XE slide scanner (Pacific Imaging, Inc.) using HistoView Software. ImageJ software (National Institutes of Health, Bethesda, MD) was used to calculate the cystic index. The images were converted to 8-bit grey scale. Equal sized non-overlapping areas were cropped covering the entire kidney image. The image threshold was adjusted similarly, and the percentage of black area was analyzed in each cropped image. Data from all cropped areas from the kidney were averaged and finally subtracted from 100 to give the cystic index as a percent.

### Tandem Affinity Purification and Immunoblotting

IMCD3 FlpIn cells stably expressing LAP tag or LAP tagged Adenylyl cyclases were lysed in buffer containing 50 mM Tris-HCl, pH 7.4, 200 mM KCl, 1 mM MgCl2, 1mM EGTA, 10 % glycerol, 1 mM DTT, 1% digitonin, 0.05% n-Dodecyl-β-D-Maltoside, 0.25% Cholesteryl hemisuccinate, 1 mM of AEBSF, 0.01 mg/mL of Leupeptin, pepstatin and chymostatin ^79^. Lysates were centrifuged at 12000xg for 10 min followed by tandem IPs ^80^. Briefly, the GFP immunoprecipitates were first digested with TEV (N terminal LAP) or PreScission (for C terminal LAP) protease for 16h at 4°C. The supernatants were subjected to secondary IPs with S-protein agarose. Treatment with Endo H and PNGase F (NEB) was performed on the IP-ed proteins on S-protein agarose beads in NEB designated buffers at 37°C for 2 h. The resulting secondary IPs were eluted in 2× urea sample buffer (4 M urea, 4% SDS, 100 mM Tris, pH 6.8, 0.2% bromophenol blue, 20% glycerol, and 200 mM DTT) at 37°C for 30 min and analyzed by immunoblotting^79^. For detection of different glycosylation forms of the LAP-tagged Adenylyl cyclases by S-tag immunoblotting and based on the stable expression levels, we required ∼0.75 ml packed cell pellet for finally eluting secondary IPs in 30-40 μl of 2× urea sample buffer from 30-40 μl S-protein agarose beads. Tandem-IPs were mostly run on 4–20% Mini-PROTEAN TGX Precast Protein Gels (Bio-Rad). Immunoblots from tandem affinity purifications were probed with antibodies against S-tag (mouse monoclonal MAC112; EMD Millipore), ANKMY2 (HPA067100, Sigma), α-tubulin (clone DM1A, T6199, Sigma), HA tag (clone 3F10, Roche) followed by visualization using IRDye-tagged secondary antibodies. The images were acquired with the Odyssey Fc Imaging System (LI-COR Biosciences), and the analysis and quantification of individual western blot bands was performed with Image Studio Lite Western Blot Analysis software (LI-COR Biosciences).

### Immunoblotting kidney samples

Frozen whole kidney tissues were homogenized using Fisherbrand™ Pellet Pestles in a RIPA lysis buffer containing 10 mM sodium phosphate (pH 7.5), 150 mM NaCl, 1.5 mM MgCl2, 0.5 mM DTT, 1% Triton X-100, 10 μg/ml each of Leupeptin, Pepstatin and Chymostatin and 0.1 mM AEBSF), Phosphatase inhibitor cocktail 2 (Sigma-Aldrich, P5726) and Phosphatase inhibitor cocktail 3 (Sigma-Aldrich, P0044). The lysates were centrifuged at 10,000 × g for 10 min at 4°C. Protein concentration was determined by bicinchoninic acid assay (Thermo Fisher Scientific). Equal amount of samples was prepared with SDS buffer containing β-mercaptoethanol, heated at 95°C for 5 min and 30 µg proteins were loaded each wall on Mini-PROTEAN TGX 4–15% gels (BIO-RAD Laboratories, Inc.). Next, proteins were blotted on polyvinylidene difluoride (PVDF) membranes. Membranes were blocked for 1 h at room temperature with TBS-Tween (0.1%) containing 1% Gelatin (Sigma), followed by incubation with the primary antibody diluted in TBS-T for overnight at 4 °C. Primary antibodies used were mouse anti-ERK (9107S; Cell Signaling Technology, 1:1,000), rabbit anti-pERK (4370S; Cell Signaling Technology, 1:1,000) and mouse anti α-tubulin (clone DM1A, Sigma; T6199) 1:5,000). Next day, the membranes were washed 5 times with TBS-T and followed by visualization using IRDye-tagged secondary antibodies and hFAB Rhodamine Anti-b-Tubulin (Bio-Rad; 12004166) for counterprobing anti-ERK blots. Images were taken in a BioRad Chemidoc MP imaging system. Densitometry was performed using default settings of the Bio-Rad Image lab software.

### Total cAMP measurements from kidneys

Frozen kidney tissue (∼100 mg) was homogenized using 1.4 mm Zirconium-Silicate spheres (Lysing Matrix D beads, 1169130-CF, MPBio) in 1 ml of 0.1 M HCl on an Omni Bead Ruptor Elite bead mill homogenizer (Revvity). After homogenization, the sample was diluted 5-fold. cAMP levels were measured using the Enzo Direct cAMP ELISA kit (ADI-900-066A) following the manufacturer’s protocol. Briefly, 50 µl of neutralizing reagent, 100 µl of sample, 50 µL of blue conjugate, and 50 µl of yellow antibody were added to each well of the ELISA plate. The plate was incubated at room temperature for 2 h. After incubation, the wells were washed three times with 400 µl of wash buffer. Then, 200 µl of substrate solution was added to each well and incubated for 1.5 h. Finally, 50 µl of stop solution was added, and absorbance was measured at 405 nm. Protein concentration was determined by bicinchoninic acid assay (Thermo Fisher Scientific). To normalize for protein content, the resulting picomole per mL determinations (pmol/ml) was divided by the total protein concentration (mg/ml) in each sample and expressed as pmol cAMP/mg protein.

### BUN measurements

Mice were euthanized and blood samples were obtained by ventricular puncture. BUN measurements were obtained using the Vitros 250 chemistry analyzer (GMI Inc., Ramsey, MN).

## QUANTIFICATION AND STATISTICAL ANALYSIS

Cystic area and length of cilia in each mouse genotype was measured using ImageJ software (NIH, Bethesda, MD). For measuring cilia positive for a particular protein, we first identified cilia using acetylated tubulin staining as a ciliary marker. In experiments where LAP-tagged constructs were used to rescue ciliary localization of proteins, we counted cilia only from GFP expressing cells. Next, we carefully counted cilia for the presence of staining corresponding to the protein of interest. All Z-planes containing acetylated tubulin staining were analyzed for staining. We did not use any threshold intensity while counting and all cilia including those showing low intensity staining were regarded as positives. All data in Figures are expressed as mean ± SD or SEM. To assess the statistical significance of differences among genotypes we performed one-way ANOVA with Sidak’s multiple comparisons tests, or unpaired, two-sided, student’s *t* tests that assumed unequal variances in treatments, as mentioned in legends. Microsoft Excel and GraphPad Prism (GraphPad, La Jolla, CA) were used for statistical analysis. Values of P<0.05 were considered significant.

## Acknowledgments

We thank the transgenic core, molecular pathology, and mouse animal care facility in UT Southwestern. We thank John Shelton for histopathology core support.

## Author contributions

Conceptualization: SM, SHH, FQ Data curation: SHH

Formal analysis: SM Funding acquisition: SM

Investigation: SHH, KC, HBB, KW, YX Methodology: SHH, KC, HBB, KW, YX Supervision: SM

Visualization: SHH, KC Validation: SHH, KC Writing—original draft: SM

Writing—review & editing: SM, SHH, KC, YX, FQ

## Data sharing statement

Mice strains described will be made available to other researchers upon request. All data are available in the main text or the supplementary materials. All expression plasmids described are available from the authors.

## Supplemental Material

This article contains the following supplemental material.

**Supplemental Figure 1.**Early cystogenesis in embryonic-onset PKD is suppressed from lack of ANKMY2.

**Supplemental Figure 2.**Lack of ANKMY2 has no effect on later cystic burden in embryonic-onset PKD.

**Supplemental Figure 3.**Cystogenesis in adult-onset PKD in male mice is suppressed from lack of ANKMY2.

**Supplemental Figure 4.**Cystogenesis in adult-onset PKD in female mice is not suppressed from lack of ANKMY2.

**Supplemental Figure 5.**Surface levels of ADCY5 and ADCY6 in *Ankmy2* ko IMCD3 cells were restored upon exogenous ^HA^ANKMY2 reexpression.

**Supplemental Figure 6.**Ciliary length increase from PC1 loss in adult kidney epithelia is suppressed from ANKMY2 loss.

