## Supplemental Figure for "Kidney cystogenesis in embryonic- and adult-onset ADPKD is suppressed from lack of adenylyl cyclase targeting to cilia"

**Table of Contents**

**Supplemental Figure 1.**

Early cystogenesis in embryonic-onset PKD is suppressed from lack of ANKMY2.

**Supplemental Figure 2.**

Lack of ANKMY2 has no effect on later cystic burden in embryonic-onset PKD.

**Supplemental Figure 3.**

Cystogenesis in adult-onset PKD in male mice is suppressed from lack of ANKMY2.

**Supplemental Figure 4.**

Cystogenesis in adult-onset PKD in female mice is not suppressed from lack of ANKMY2.

**Supplemental Figure 5.**

Surface levels of ADCY5 and ADCY6 in *Ankmy2* ko IMCD3 cells were restored upon exogenous $^{HA}$ANKMY2 reexpression.

**Supplemental Figure 6.**

Ciliary length increase from PC1 loss in adult kidney epithelia is suppressed from ANKMY2 loss.

**Supplemental Figure legends**

**Supplemental Figure 1. Early cystogenesis in embryonic-onset PKD is suppressed from lack of ANKMY2.**
**(A)** Images of multiple kidneys at P3 used for analyses shown for the designated genotypes. Scale, 1 mm.
**(B)** Transcript levels of *Pkd1* and *Ankmy2* in whole kidneys at P3 shown.

**Hwang_Figure S1**

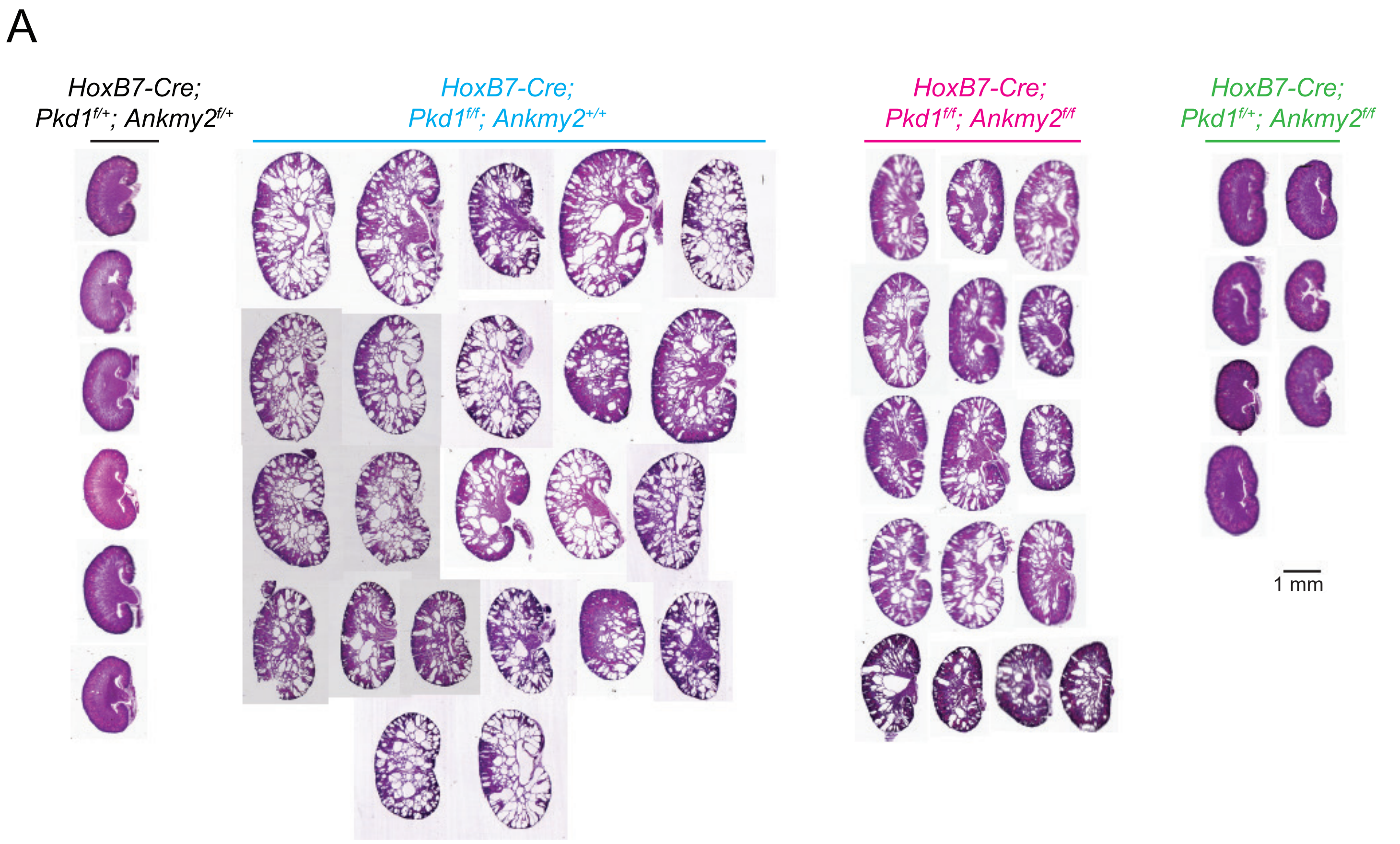


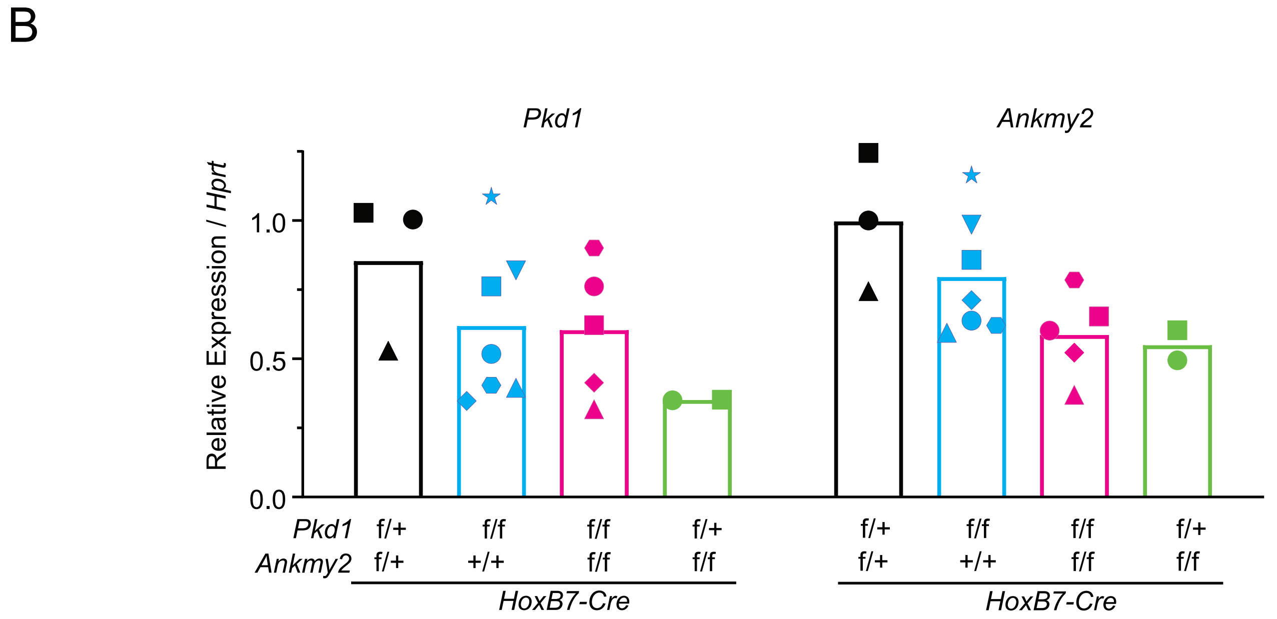

**Supplemental Figure 2. Lack of ANKMY2 has no effect on later cystic burden in embryonic-onset PKD.**
Images of multiple kidneys at P15 used for analyses shown for the designated genotypes. Scale, 2 mm.

### Hwang_Figure S2

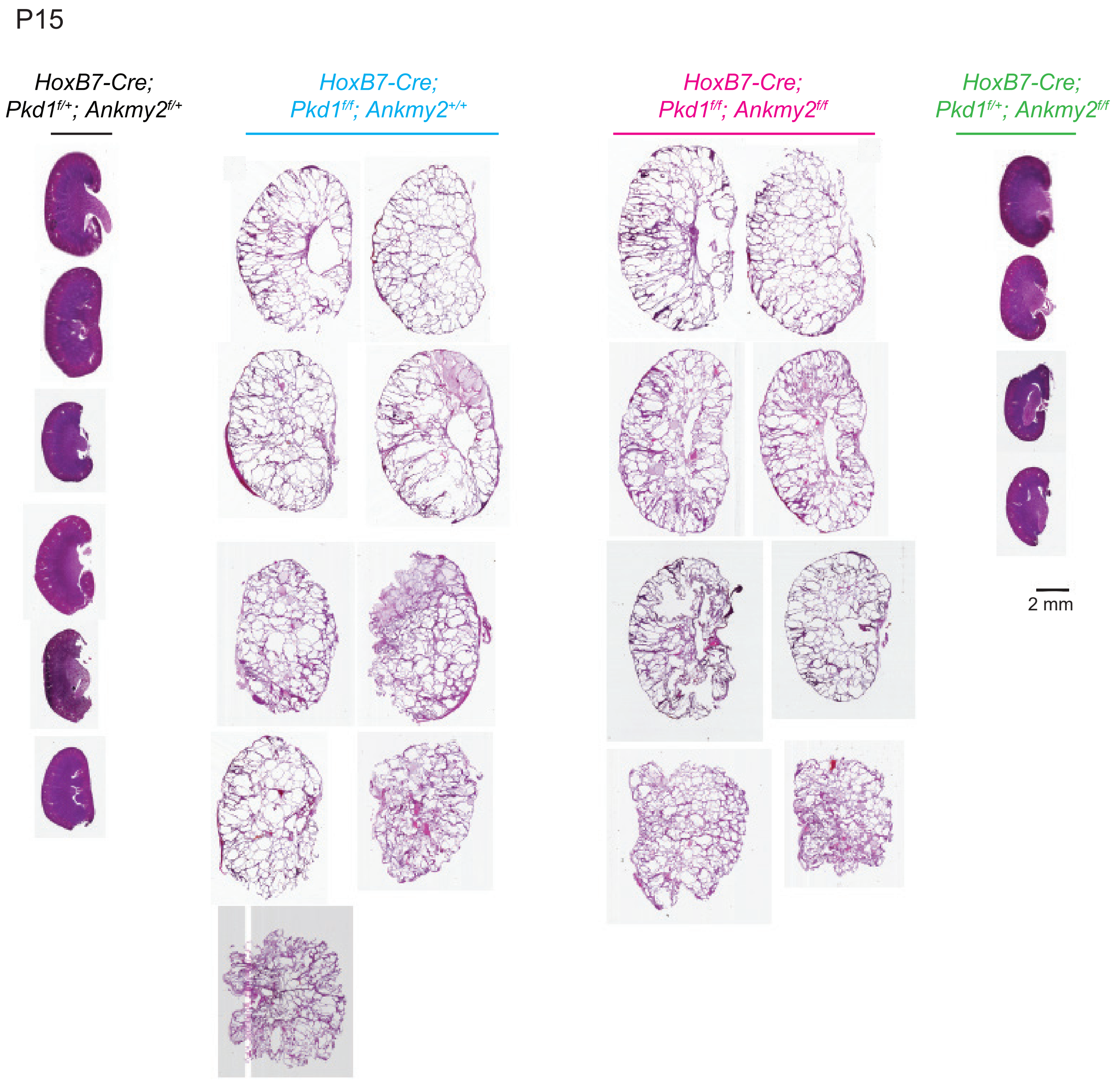

**Supplemental Figure 3. Cystogenesis in adult-onset PKD in male mice is suppressed from lack of ANKMY2.**

(**A**) qRT-PCR of whole kidneys showed depletion of *Pkd1* or *Ankmy2* in the respective inducible conditional knockout animals at 5 months. Samples are color coded to show respective levels in the same animal.

**(B)** Images of multiple kidneys from males at 5 months used for analyses shown for the designated genotypes. Scale, 2 mm.

**Hwang_Figure S3**

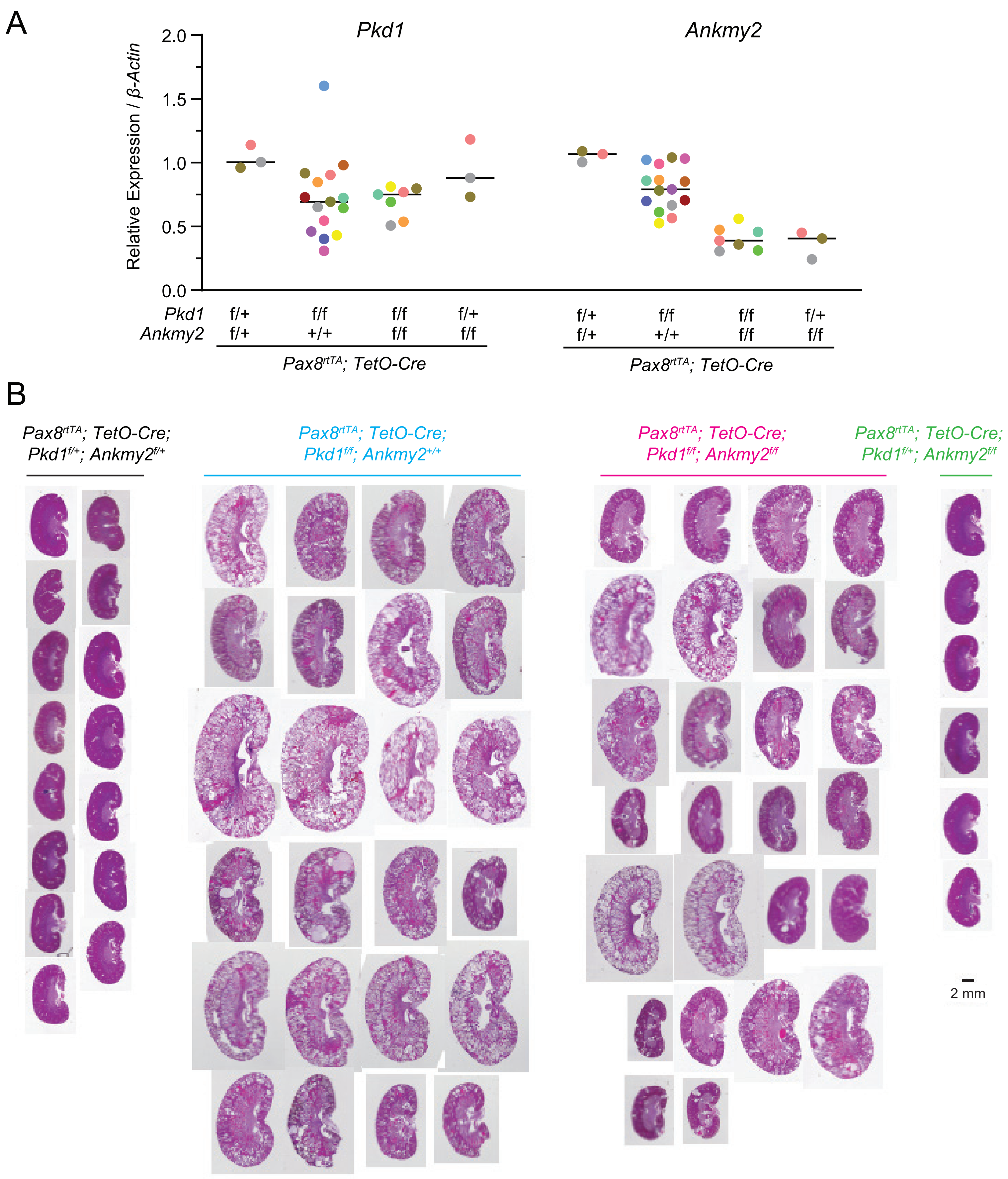

**Supplemental Figure 4. Cystogenesis in adult-onset PKD in female mice is not suppressed from lack of ANKMY2.**

**(A)** 2-Kidney to body weight ratios of 5-month-old *Pax8*$^{rtTA}$; *TetO-Cre*; *Pkd1*$^{f/f}$ female mice were not significantly different than *Pax8*$^{rtTA}$; *TetO-Cre*; *Pkd1*$^{f/f}$*; Ankmy2*$^{f/f}$.

**(B)** Cystic index in *Pax8*$^{rtTA}$; *TetO-Cre*; *Pkd1*$^{f/f\ f}$ *fe*male mice compared to *Pax8*$^{rtTA}$; *TetO-Cre*; *Pkd1*$^{f/f}$*; Ankmy2*$^{f/f}$ showed no difference.

**(C)** Cyst sizes in *Pax8*$^{rtTA}$; *TetO-Cre*; *Pkd1*$^{f/f}$ and *Pax8*$^{rtTA}$; *TetO-Cre*; *Pkd1*$^{f/f}$*; Ankmy2*$^{f/f}$ female mice were not significantly different from each other in both LTL+ and AQP2+ cysts as counted in Figure 3E-F. Superplots of N=3 kidneys/genotype are shown with different shapes and averages are plotted with larger shapes. Cystic indices in *Pax8*$^{rtTA}$; *TetO-Cre*; *Pkd1*$^{f/f}$ were 56, 44, and 42, whereas that in *Pax8*$^{rtTA}$; *TetO-Cre*; *Pkd1*$^{f/f}$*; Ankmy2*$^{f/f}$ were 39, 32, and 23.

**(D)** Images of multiple kidneys from females at 5 months used for analyses shown for the designated genotypes. Scale, 2 mm.

### Hwang_Figure S4

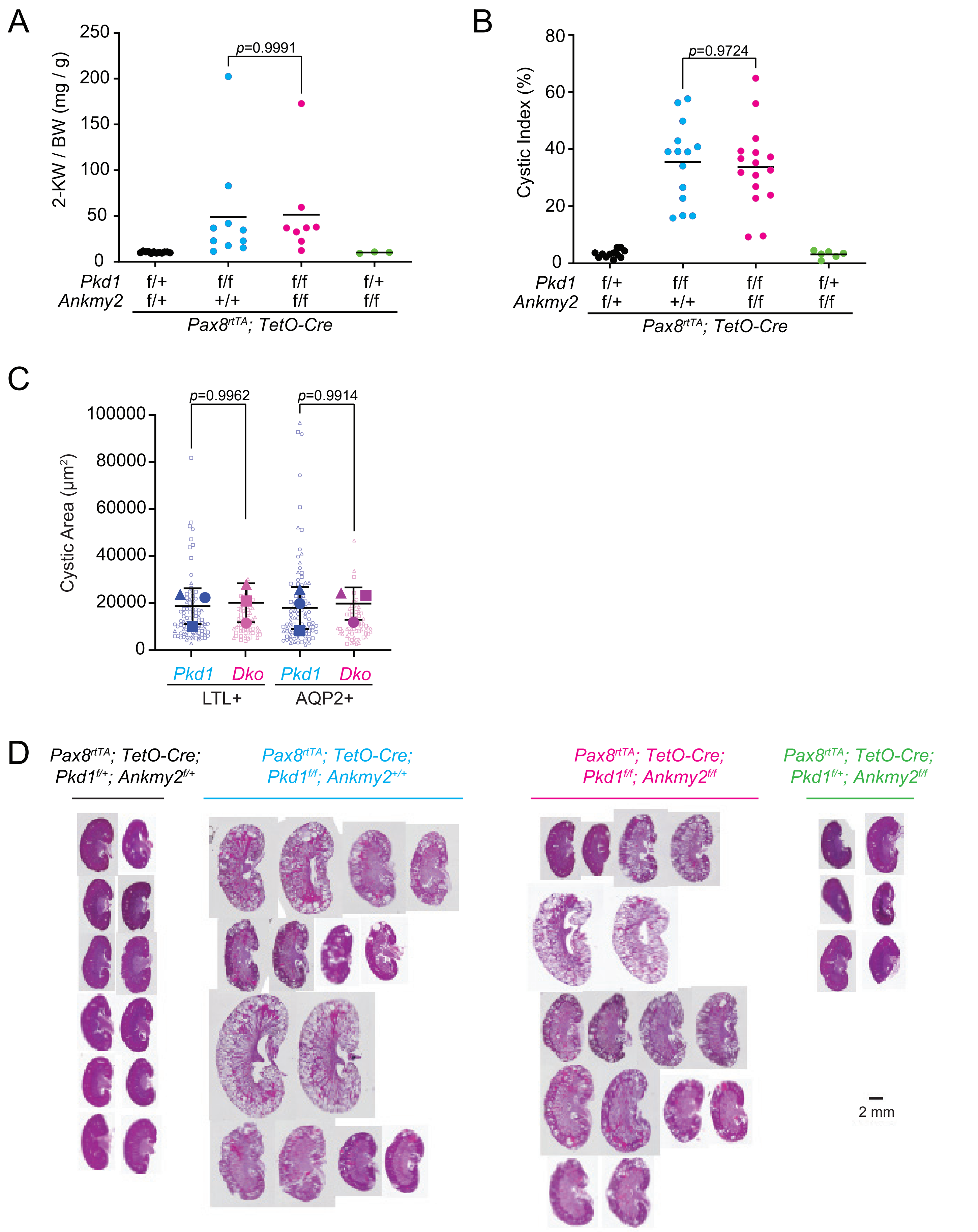

**Supplemental Figure 5. Surface levels of ADCY5 and ADCY6 in *Ankmy2* ko IMCD3 cells were restored upon exogenous $^{HA}$ANKMY2 reexpression.**

(A-B) ADCY5$^{LAP}$ and $^{LAP}$ADCY6 were localized to the plasma membrane and secretory pathway in stably expressing IMCD3 FlpIn cell line. Surface levels were reduced in *Ankmy2* ko but restored back upon $^{HA}$ANKMY2 reexpression, as detected upon performing immunostaining with antibodies against GFP, β-catenin and HA. Scale: 25 μm.

(C) Sequence of *Ankmy2* exon1 targeted for CRISPR-Cas9 mediated *Ankmy2* knockout in mouse IMCD3 Flp-In cells stably expressing ADCY5$^{LAP}$ or $^{LAP}$ADCY6. Immunoblotting showing absence of ANKMY2 in the clonal knockout lines are depicted in Figure 5G.

**Hwang_Figure S5**

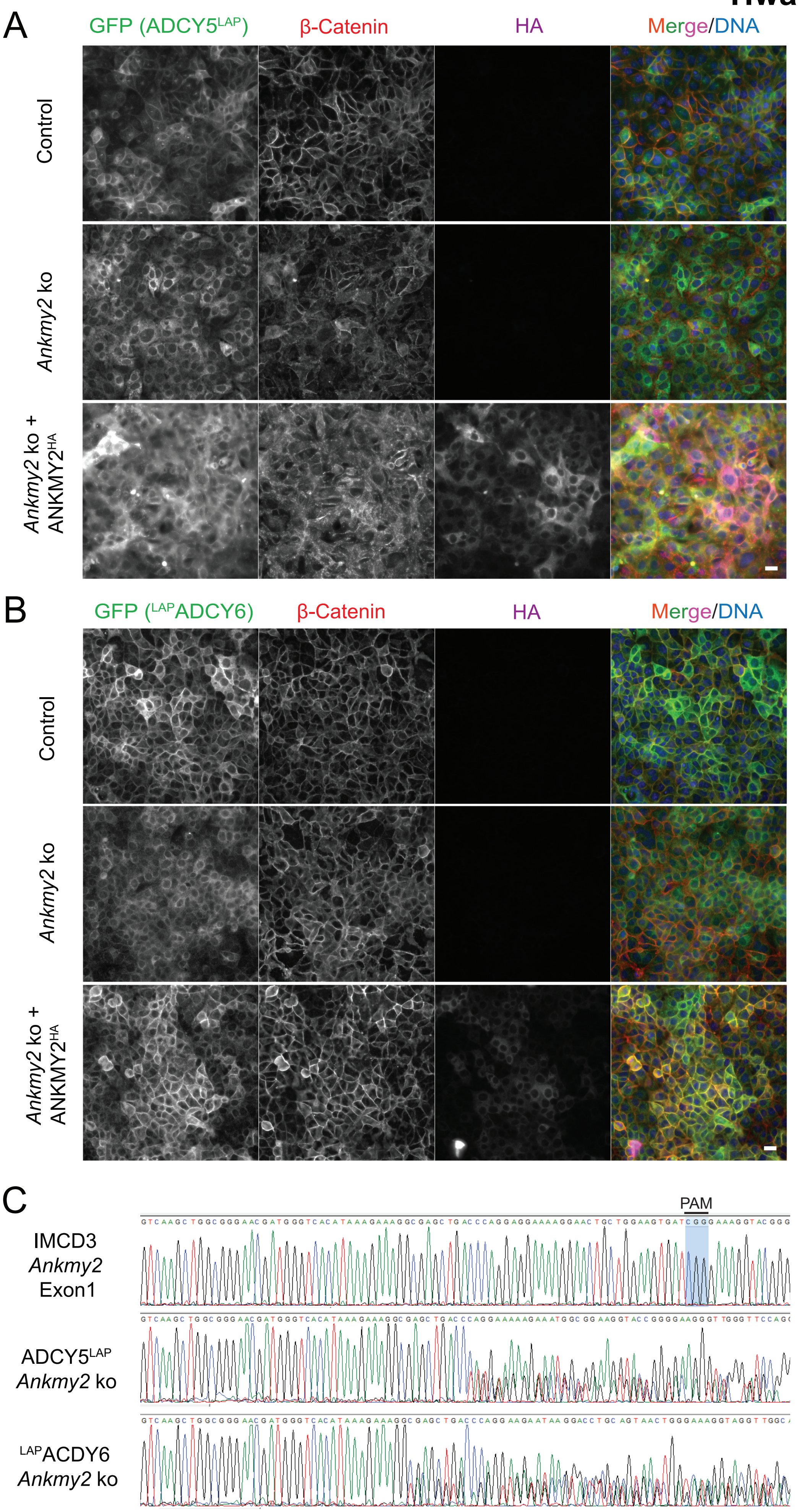

**Supplemental Figure 6. Ciliary length increase from PC1 loss in adult kidney epithelia is suppressed from ANKMY2 loss.**
**(A)** Cilia lengths were quantified and super plots of N=2-3 female 5-month-old mice shown for each genotype as in Figure 6A-B. Lengths from each kidney is shown with different shapes and averages are plotted with larger shapes.
**(B)** H&E images, 2-kidney to body weight ratios, and cystic indices of 10-week-old kidneys in control, *Pax8*$^{rtTA}$; *TetO-Cre*; *Pkd1*$^{f/f}$ and *Pax8*$^{rtTA}$; *TetO-Cre*; *Pkd1*$^{f/f}$*; Ankmy2*$^{f/f}$ male mice in C57BL/6J background shown. Kidney sections were immunostained for acetylated tubulin, AQP2 and LTL and counterstained with DAPI. Scale 10 μm.

**Hwang_Figure S6**

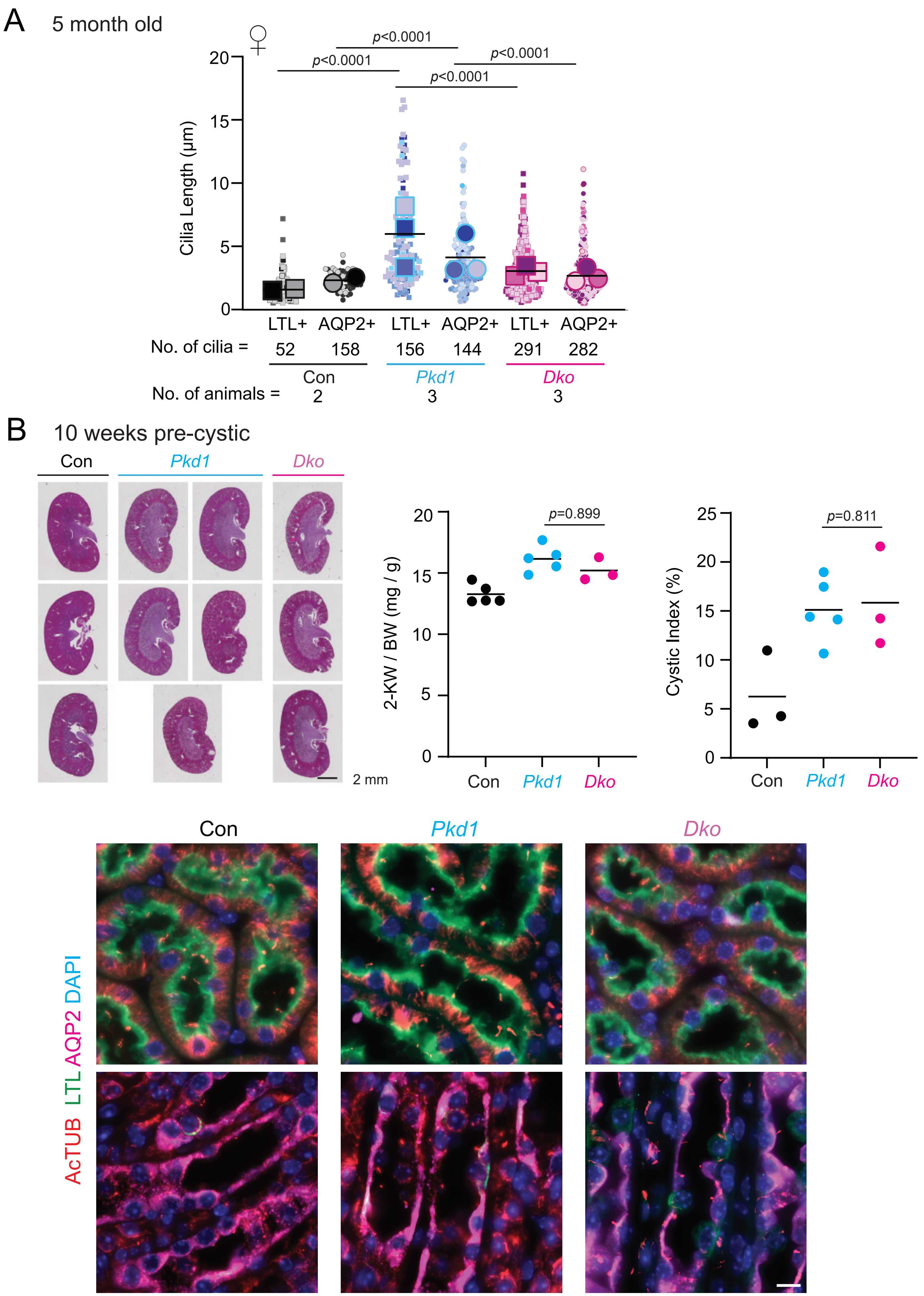
